# Regulation of bacterial surface attachment by a network of sensory transduction proteins

**DOI:** 10.1101/554154

**Authors:** Leila M. Reyes Ruiz, Aretha Fiebig, Sean Crosson

## Abstract

Bacteria are often attached to surfaces in natural ecosystems. A surface-associated lifestyle can have advantages, but shifts in the physiochemical state of the environment may result in conditions in which attachment has a negative fitness impact. Therefore, bacterial cells employ numerous mechanisms to control the transition from an unattached to a sessile state. The *Caulobacter crescentus* protein HfiA is a potent developmental inhibitor of the secreted polysaccharide adhesin known as the holdfast, which enables permanent attachment to surfaces. Multiple environmental cues influence expression of *hfiA*, but mechanisms of *hfiA* regulation remain largely undefined. Through a forward genetic selection, we have discovered a multi-gene network encoding a suite of two-component system (TCS) proteins and transcription factors that coordinately control *hfiA* transcription and surface adhesion. The hybrid HWE-family histidine kinase, SkaH, is central among these regulators and forms heteromeric complexes with the kinases, LovK and SpdS. The response regulator SpdR indirectly inhibits *hfiA* expression by activating two XRE-family transcription factors that directly bind the *hfiA* promoter to repress its transcription. This study provides evidence for a model in which a consortium of environmental sensors and transcriptional regulators integrate environmental cues at the *hfiA* promoter to control the attachment decision.

**Author summary:** Living on a surface within a community of cells confers a number of advantages to a bacterium. However, the transition from a free-living state to a surface-attached lifestyle should be tightly regulated to ensure that cells avoid adhering to toxic or resource-limited niches. Many bacteria build adhesive structures at their surfaces that enable attachment. We sought to discover genes that control development of the *Caulobacter crescentus* surface adhesin known as the holdfast. Our studies uncovered a network of signal transduction proteins that coordinately control the biosynthesis of the holdfast by regulating transcription of the holdfast inhibitor, *hfiA*. We conclude that *C. crescentus* uses a multi-component regulatory system to sense and integrate environmental information to determine whether to attach to a surface, or to remain in an unattached state.

## Introduction

In natural ecosystems, bacteria often live in surface attached, multicellular communities known as biofilms [1]. Biofilms enhance resistance to toxic compounds, promote sorption of nutrients, and facilitate exchange of genes and gene products [2]. In aquatic systems, metabolizable substrates accumulate at surfaces [3], and thus attachment can provide a fitness advantage [4]. However, it is important for cells to be able to disperse in cases when surfaces accumulate toxins, or become depleted of substrates or electron acceptors. It is therefore not surprising that surface attachment is a highly regulated process in bacteria.

The alphaproteobacterium *Caulobacter crescentus* exhibits a dimorphic developmental cycle that results in two cell types after division: a flagellated swarmer cell and a sessile stalked cell [5] (Fig 1A). The stalked cell is the reproductive cell type and divides asymmetrically, giving birth to the non-reproductive swarmer cell [5]. Swarmer cells differentiate into stalked cells, whereupon they can secrete a unipolar polysaccharide adhesin known as the holdfast [5–8] (Fig 1A). Holdfast mediated adherence to solid surfaces is permanent [9–11]; the holdfast enables cell attachment to a variety of chemically diverse materials [12], and facilitates partitioning of cells to the air-liquid interface in aqueous medium [13]. *C. crescentus* tightly regulates holdfast synthesis and thus the transition between a planktonic and a surface-attached lifestyle, which likely reduces the probability of becoming restricted to a sub-optimal environment.

**Fig 1.**
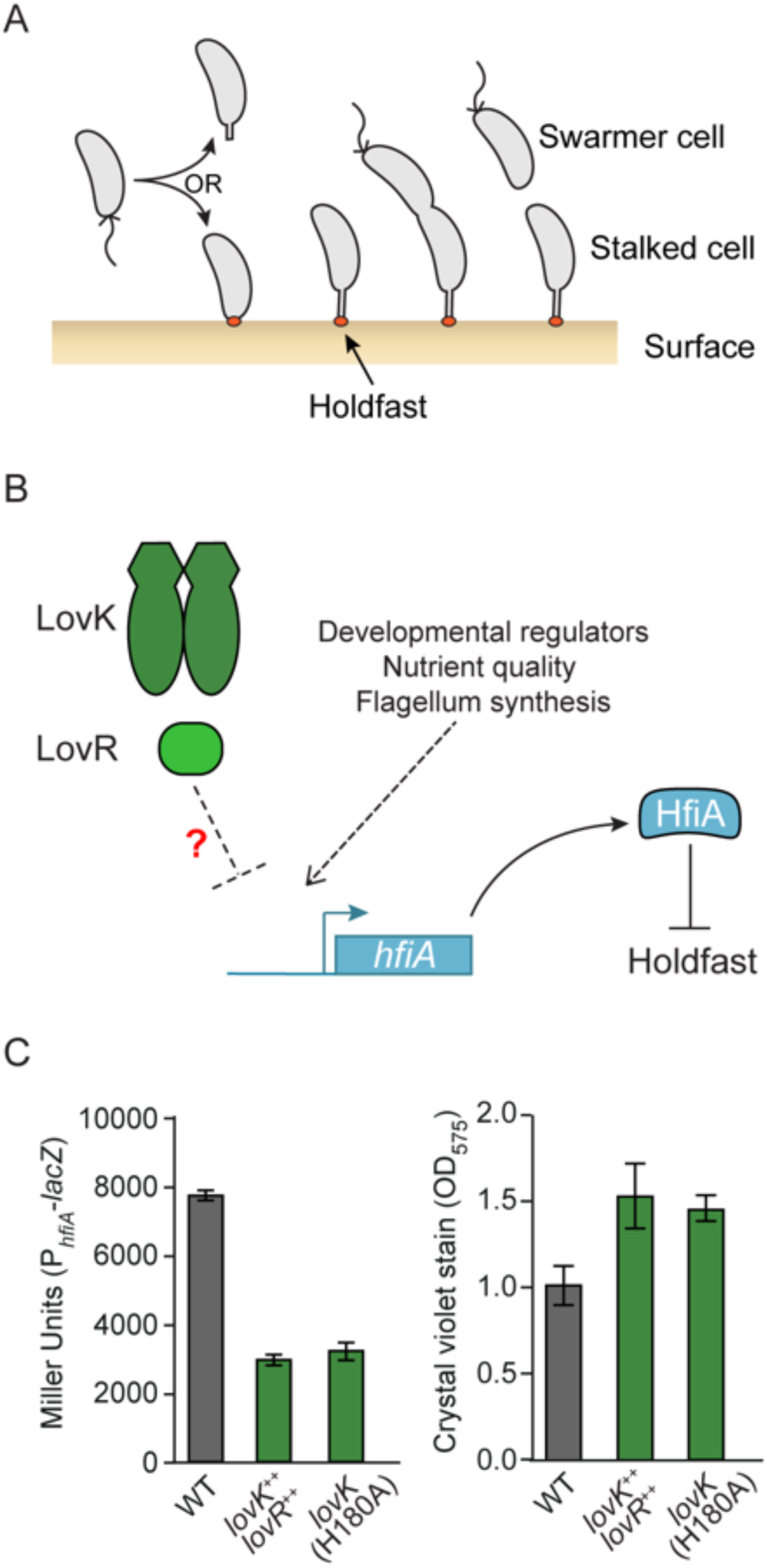
*lovK/lovR* regulation of *hfiA* expression and surface adhesion. **A)** The dimorphic bacterium *C. crescentus* builds a polysaccharide-based adhesin known as the holdfast (orange) at one cell pole, which mediates irreversible surface attachment. **B)** Transcription of the holdfast inhibitor, *hfiA*, is regulated by developmental factors [14], flagellum synthesis [15], and environmental response systems [14] including the LovK/LovR two-component system [14]. The goal of this study is to understand the mechanism of *hfiA* transcriptional regulation by LovK/LovR. **C)** Characterization of *hfiA* expression and surface attachment in wild-type (WT), *lovK/lovR* overexpression (*lovK*^++^*lovR*^++^) and *lovK*(H180A) strains. Left: **β**-galactosidase activity in strains carrying a P*_hfiA_*-*lacZ* transcriptional fusion. Right: Crystal violet staining of surface attached cells after growth in polystyrene wells. Strains were grown in defined M2X medium. Bars represent mean ± s.d.; n=6 for **β**-galactosidase assay and n=8 for crystal violet stain.

The small protein HfiA is a potent inhibitor of holdfast synthesis [14]. HfiA directly targets HfsJ, a glycolipid glycosyltransferase required to build the holdfast [14]. The promoter of *hfiA* can integrate a number of regulatory cues that modulate its transcription and thereby affect holdfast production (Fig 1B). Multiple cell cycle regulators bind and regulate the *hfiA* promoter [14]. In addition, flagellar assembly indirectly affects *hfiA* transcription by an unknown mechanism [15]. Environmental signals such as nutrient quality and the two-component system (TCS) proteins LovK-LovR [14] and MrrA [16] influence *hfiA* transcription by unknown and independent mechanisms. CleA, a CheY-like response regulator that tunes flagellar rotation in response to cyclic-di-GMP binding, also regulates holdfast synthesis, though it is not known if CleA-mediated holdfast regulation involves HfiA [17]. In an effort to better understand the complex regulatory network that controls expression of the holdfast inhibitor, HfiA, we have investigated the molecular basis by which the LovK-LovR TCS controls *hfiA* transcription.

TCS are commonly used by bacteria to sense environmental signals and regulate adaptive changes in cell physiology [18]. Archetypal TCS consist of a sensor histidine kinase (HK) and a DNA-binding response regulator (RR), which generally form simple linear pathways that couple environmental perturbation to a change in gene expression. There is good experimental support for the paradigm that TCS form insulated signaling systems, with little cross-regulation between non-cognate kinases and responses regulators [19–22], though there are examples of branched TCS pathways in which sensor kinases signal to more than one regulator, or in which signals from multiple kinases converge on a single regulator (reviewed in [23, 24]). Complex cellular processes including sporulation [25], stress responses [16, 26–29], biofilm formation [30], and cell cycle [31, 32] are often controlled by multiple TCS in branched pathways. In this study, we have defined a regulatory system composed of multiple TCS proteins and transcription factors that regulates holdfast development in *C. crescentus*.

We have previously shown that the TCS LovK-LovR regulates holdfast development [14] and surface adhesion [14, 33] by influencing the transcription of *hfiA* [14]. LovK is a soluble photosensory HK that contains a LOV (light, oxygen, voltage) sensor domain [34] N-terminal to a HWE/HisKA_2 family [35, 36] kinase domain. The LOV domain binds a flavin mononucleotide (FMN) cofactor [37], which confers the ability to sense blue light and the redox state of the environment [33, 37]. LovK is encoded adjacent to LovR, a single domain response regulator (SDRR). The regulation of *hfiA* transcription by LovK-LovR must be indirect as LovR has no DNA-binding domain. We developed a genetic selection aimed at identifying genes that function together with or downstream of LovK-LovR to control *hfiA* transcription. We found that the hybrid histidine kinase SkaH and the canonical TCS SpdS-SpdR function with LovK-LovR to control transcription of *hfiA*. LovK and SpdS form heteromeric complexes with SkaH *in vitro* and *in vivo*, providing evidence for cross-regulation between these different TCS. We further demonstrate that the response regulator SpdR indirectly regulates *hfiA* transcription by activating expression of two direct transcriptional repressors of *hfiA*, RtrA and RtrB. Our data provide evidence for a complex regulatory system that has the capacity to regulate *hfiA* expression and surface attachment in response to multiple environmental signals.

## Results

### A *lovK*(H180A) mutant phenocopies a *lovK-lovR* overexpression strain

Coordinate overexpression of *lovK*-*lovR* increases the probability that any single cell will develop a holdfast, and therefore enhances adhesion of a cell population to surfaces [14, 33]. This phenotype is due to the repressive effect of *lovK*-*lovR* expression on *hfiA* transcription [14]. During the course of our studies, we discovered that mutating the conserved histidine phosphorylation site of LovK to alanine - *lovK*(H180A) - resulted in decreased *hfiA* transcription and a concomitant increase in surface attachment (Fig 1C). In other words, a strain in which the chromosomal copy of *lovK* is replaced with *lovK*(H180A) phenocopies a *lovK-lovR* overexpression strain. To further investigate the role of the LovK-LovR TCS proteins, we ectopically complemented a Δ*lovK-lovR* strain with either the wild-type allele of *lovK* or the *lovK*(H180A) allele. In the absence of *lovR*, expression of the wild-type *lovK* allele had no effect on surface attachment. On the other hand, expression of the *lovK*(H180A) allele increased surface adhesion (Fig S1). These results indicate that expressing the unphosphorylatable *lovK*(H180A) allele is sufficient to enhance surface attachment, and that *lovR* is not required for this phenotype. We chose to use the strain with the *lovK*(H180A) allele expressed from its native locus to investigate the molecular connection between the LovK sensor kinase and *hfiA* transcription as it *1)* provided a cleaner and more stable genetic background than the *lovK-lovR* plasmid overexpression strain, *2)* expressed *lovK*(H180A) from the native *lovK* promoter, and *3)* did not require the addition of metabolizable inducers.

### A forward genetic selection identifies genes that function with *lovK-lovR* to repress *hfiA* **expression**

The SDRR LovR lacks a DNA-binding output domain, and thus transcriptional regulation of *hfiA* by LovK-LovR is likely indirect. Moreover, LovR is not required to regulate *hfiA* transcription when LovK lacks its histidine phosphorylation site. We hypothesized that additional regulator(s) function either downstream or in concert with LovK-LovR to repress *hfiA* transcription. To identify such regulator(s), we designed a forward genetic selection to isolate mutants in which the repressive regulatory connection between LovK-LovR and *hfiA* was disrupted (Fig 2A). Specifically, we fused the hfiA promoter (P*_hfiA_*) to the chloramphenicol acetyltransferase (cat) gene and transformed the lovK(H180A) strain with this reporter plasmid (P*_hfiA_-CAT*). We then used growth of this strain in the presence of chloramphenicol as a proxy for P*_hfiA_* activity. By serially passaging this reporter strain, we selected for spontaneous mutants with enhanced growth in the presence of chloramphenicol (Fig 2A). We predicted that this approach would enable us to identify mutants in which genetic connection between *lovK-lovR* and *hfiA* transcription was disrupted.

**Fig 2.**
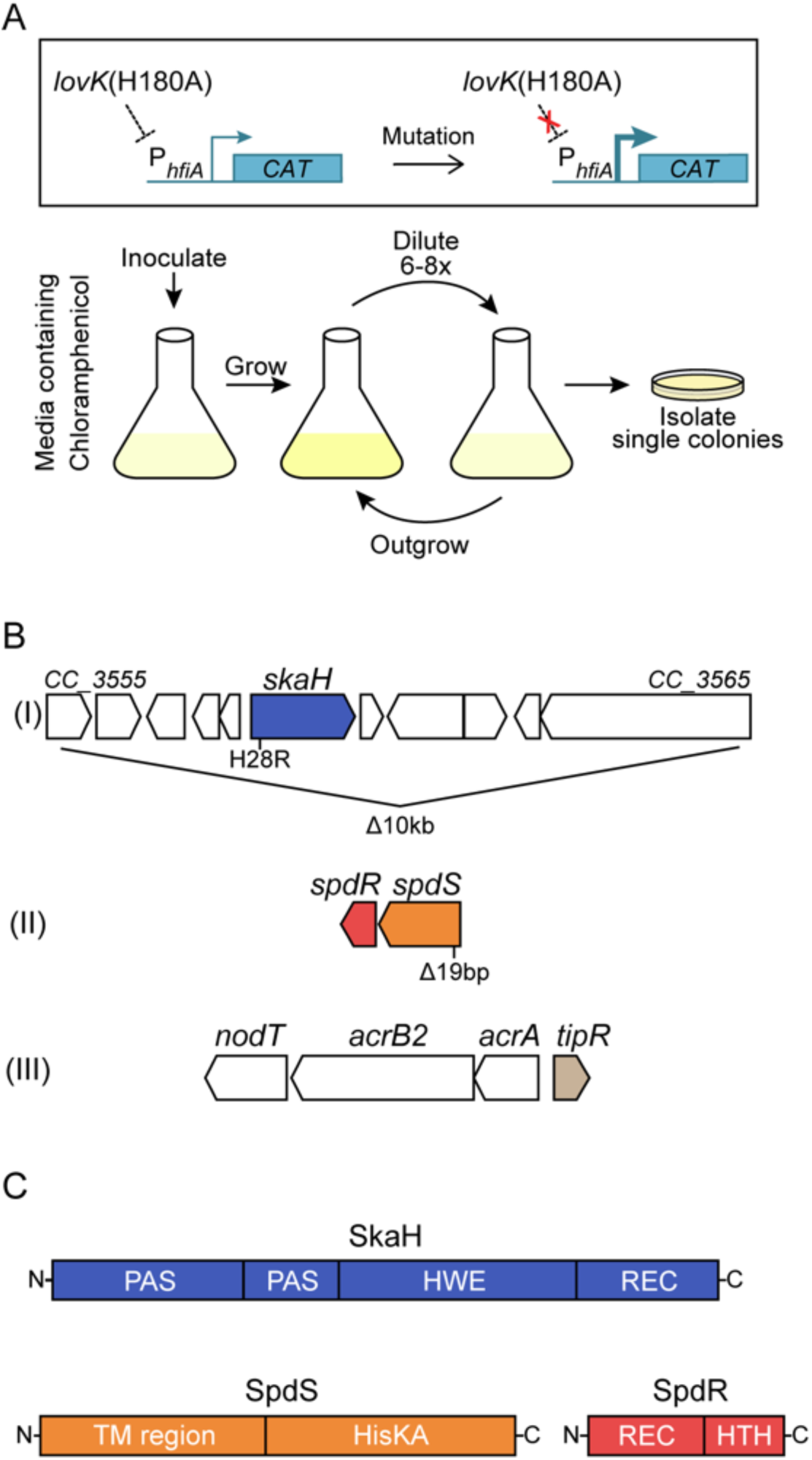
Genetic selection identifies regulators that function with *lovK*(H180A) to repress *hfiA*. **A)** A strategy to select for spontaneous mutants that disrupt *lovK*(H180A) repression of the *hfiA* promoter. The P*_hfiA_-CAT* transcriptional fusion allows for selection of mutants with increased hfiA promoter activity. Specifically, cultivation in M2X medium supplemented with chloramphenicol allows for selection of enhanced expression of chloramphenicol acetyltransferase (CAT) from P*_hfiA_-CAT*. B) Mutations in three genomic regions identified in the selection with specific lesions marked below the genetic loci. Lesions at the *acr* locus confer increased chloramphenicol resistance irrespective of *lovK* (Fig S3); the mutant alleles identified at this locus are not indicated in the diagram but are listed in Table S1. **C)** Predicted domain organization of the TCS proteins SkaH, SpdS and SpdR. PAS – sensory domain, HWE and HisKA – histidine kinase domains, REC – receiver domain, TM – transmembrane, HTH – helix-turn-helix DNA-binding domain.

After several passages, we isolated dozens of mutants that grew faster than the parent strain. In these isolates, we first sequenced the P*_hfiA_-CAT* promoter fragment to identify clones with cis mutations in the hfiA promoter itself that increase promoter activity. In mutants with an intact P*_hfiA_-CAT* promoter fragment, we assessed P*_hfiA_* activity in two independent secondary screens; we measured activity of a second hfiA promoter fusion (P*_hfiA_-lacZ*), and assessed activity of the native hfiA promoter by measuring the bulk adhesion phenotype of these mutants. Two classes of mutants emerged from this selection: 1) strains that grew faster in chloramphenicol as a result of P*_hfiA_-CAT* promoter mutations, but that displayed unaltered surface adhesion and no change in P*_hfiA_-lacZ* activity compared to the parental lovK(H180A) strain (Fig S2), and 2) mutants with an intact P*_hfiA_-CAT* reporter plasmid that displayed fast growth in chloramphenicol, reduced surface adhesion, and increased β-galactosidase activity from the P*_hfiA_-lacZ* plasmid (Table S1). We mapped polymorphisms in nine spontaneous mutants from this latter class, derived from three independent selections, by whole genome sequencing.

Two mutants derived from the same flask shared a ~10kb deletion that included gene *CC_3560* (*CCNA_03675*), annotated as a hybrid histidine kinase (Fig 2B and Table S1). A single mutant had a 19bp deletion that resulted in a frameshift in the transmembrane histidine kinase gene *spdS* (Fig 2B and Table S1). The remaining six mutants had lesions in either *tipR* or both *acrB2* and *tipR* (Fig 2B and Table S1). TipR is a repressor of the *acrAB2-nodT* operon, which encodes the components of the AcrAB2-NodT antibiotic efflux pump [38]. As several of our mutants harbored lesions in *tipR*, we hypothesized that enhanced growth in chloramphenicol in these strains was due to de-repressed expression of the efflux pump. Deletion of *tipR* in either wild-type or the *lovK*(H180A) background increased growth rate in the presence of chloramphenicol (Fig S3). We concluded that disruption of *tipR* is a general genetic solution to combat chloramphenicol toxicity, and that these mutants were confounding the efficiency of our selection. We therefore modified our selection strategy and ectopically expressed a second copy of *tipR* from its native promoter in an effort to minimize enrichment of *tipR* mutants. Using this strategy, we identified nine mutants from three independent selections with a point mutation resulting in a *CC_3560*(H82R) allele. Altogether, our results provided evidence that the sensor kinases CC_3560 and SpdS play a role in *hfiA* repression by LovK-LovR. Though we likely did not saturate our selection, we decided to move forward with our initial hits of the TCS proteins CC_3560 and SpdS-SpdR.

### *CC_3560* (*skaH*) and *spdR* regulate *hfiA* transcription and adhesion downstream of *lovK*

Our selection identified mutations in two sensor kinase genes: *CC_3560* and *spdS. CC_3560* encodes a cytoplasmic hybrid histidine kinase (Fig 2C) that, like LovK, belongs to the non-canonical HWE/HisKA_2 family of histidine kinases [36]. We have named this kinase *skaH* (*<u>s</u>*ensor *<u>k</u>*inase *<u>a</u>*ssociated *<u>h</u>*ybrid) for reasons we describe in a later section. *spdS* is the transmembrane sensor kinase component of the SpdS-SpdR TCS, which has been reported to regulate genes involved in stationary phase adaptation in *C. crescentus* [39, 40] (Fig 2C). Homologs of SpdS-SpdR in other species are more commonly known as RegB-RegA, and have been reported to detect changes in cellular redox state, and regulate a variety of energy generating and energy consuming processes (Reviewed in [41]).

To validate the results of our genetic selection, we tested whether the TCS genes *skaH* and *spdS* are involved in regulation of *hfiA* transcription and surface adhesion. Though mutations in *spdR* were not identified in our selection, we also tested its role in *hfiA* regulation and adhesion since it is the cognate RR of SpdS. We first generated in-frame deletions of *skaH, spdS* or *spdR* in the *lovK*(H180A) genetic background to determine if any of these genes were required for LovK(H180A) to modulate *hfiA* and adhesion. Repression of *hfiA* transcription by *lovK*(H180A) required both *skaH* and *spdR* (Fig 3A). Similarly, deletion of *skaH* or *spdR* attenuated the hyperadhesive phenotype of *lovK*(H180A) (Fig 3B). Deleting *spdS* also ablated repression of *hfiA* by *lovK*(H180A) (Fig 3A). We note that the surface adhesion phenotype of the *lovK*(H180A)D*spdS* strain varied depending on the growth phase of the culture. At early time points, surface adhesion of *lovK*(H180A)Δ*spdS* cultures is lower than *lovK*(H180A) cultures. When cultures begin to enter stationary phase, we observed a sharp increase in surface adhesion in the *lovK*(H180A)Δ*spdS* cultures that eventually matched the levels we observed in *lovK*(H180A) (Fig 3B). Importantly, the *skaH, spdR* and *spdS* deletion phenotypes we observed on the *lovK*(H180A) background were complemented by ectopic expression of these deleted genes from a xylose-inducible promoter (Fig S4). Together, these data provide evidence that *skaH and spdR* are required for *lovK*(H180A)-dependent regulation of *hfiA* transcription and surface adhesion. *lovK*(H180A)-dependent regulation of *hfiA* transcription and adhesion requires *spdS* at low culture density.

**Fig 3.**
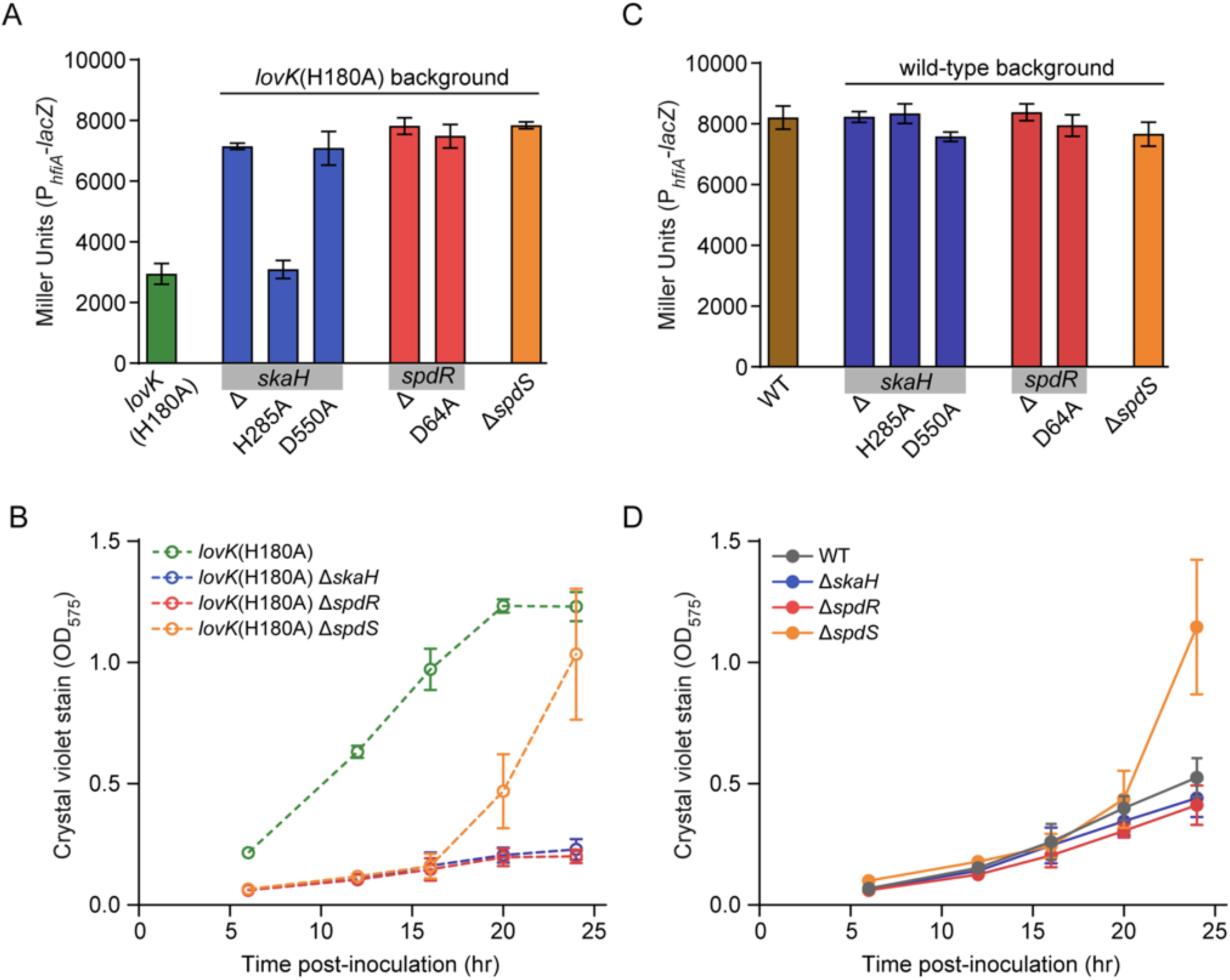
TCS genes identified in the selection are necessary for *lovK*(H180A) to repress *hfiA* transcription. **A and C**) P*_hfiA_-lacZ* activity in strains bearing in-frame deletions (Δ) or the indicated substitutions of the conserved phosphorylation sites of *skaH, spdS* and *spdR* in a *lovK*(H180A) background (A) and in a wild-type background (C). Bars represent mean ± s.d.; n=5. **B and D**) Attachment of cells to polystyrene plates measured by crystal violet stain. Strains bear in-frame deletions of *skaH, spdS* or *spdR* in a *lovK*(H180A) background (B) and in a wild-type background (D). Bars represent mean ± s.d.; n=6 for each strain at each timepoint.

We next asked whether this set of TCS genes affect *hfiA* transcription and surface adhesion in wild-type cells. Activity from the P*_hfiA_-lacZ* transcriptional reporter in the ΔskaH and ΔspdR strains was indistinguishable from wild type (Fig 3C). We note that under these conditions, transcription of *hfiA* is high and that further increases are difficult to measure. Similarly, under these conditions wild-type surface attachment is low (Fig 3D). Nevertheless, we detect a reproducible and statistically significant decrease in adhesion of strains carrying deletions of *skaH* or *spdR* compared to wild type (Fig 3D and Fig S5). Deletion of *spdS* in a wild-type background resulted in a similar adhesion phenotype to the *lovK*(H180A)Δ*spdS* strain, where we observed a marked increase in adhesion relative to wild type as the culture entered stationary phase (Fig 3D). We note that the Δ*spdS* hyperattachment phenotype during stationary phase depends on holdfast synthesis (Fig S6). These data provide evidence that under these growth conditions, SkaH and SpdR play subtle roles in regulation of adhesion. The role of SpdS depends strongly on growth phase.

### Conserved SkaH and SpdR phosphorylation sites are required for *hfiA* transcriptional regulation

We next addressed the role of the conserved *skaH, spdS,* and *spdR* phosphorylation sites in regulation of *hfiA* transcription by *lovK*(H180A). The hybrid histidine kinase SkaH has a histidine phosphorylation site in its kinase domain and an aspartyl phosphorylation site in its receiver (REC) domain. We individually mutated these chromosomal codons to encode alanine (resulting in H285A and D550A alleles, respectively) in a *lovK*(H180A) background. The *skaH*(H285A) mutation had no effect on *lovK*(H180A) repression of *hfiA* transcription (Fig 3A). The *skaH*(D550A) mutation ablated *hfiA* repression by *lovK*(H180A) (Fig 3A) and ablated its hyperadhesive phenotype (Fig S7) similar to the *ΔskaH* mutation. Mutating the aspartyl phosphorylation site of the response regulator, SpdR, to alanine (D64A) also blocked *hfiA* repression by *lovK*(H180A) (Fig 3A) and prevented hyperadhesion (Fig S7) similar to the *ΔspdR* mutation. Similar to the null alleles, the H®A or D®A phosphosite mutations in *skaH* and *spdR* in a wild-type background had no effect on *hfiA* promoter activity (Fig 3C), and a minor effect on surface adhesion (Fig S7). The transcription and adhesion phenotypes of *spdS*(H248A) were highly variable from day to day for reasons we do not yet understand, and thus we cannot presently draw any conclusions about this strain. Overall, we conclude that intact phosphorylation sites on the *skaH* and *spdR* REC domains are required for *lovK*(H180A) dependent regulation of *hfiA* transcription and surface adhesion.

### SkaH directly interacts with LovK and SpdS *in vitro* and *in vivo*

In an effort to understand the molecular basis of these genetic interactions between TCS genes, we tested whether the corresponding TCS proteins physically interact. We first performed a bacterial two-hybrid (B2H) assay in a heterologous *E. coli* system [42], fusing the LovK, SkaH or SpdS kinases to the C-terminus of the T25 and T18c split domain fragments of adenylate cyclase. We note that the transmembrane domain was not included in the SpdS construct in these experiments, which contains only the cytoplasmic kinase domain (hereafter named SpdS*). Sensor histidine kinases generally form stable homodimers [43–45]. Co-expression of the same kinase fused to the T25 and T18c fragments resulted in blue colonies on X-gal medium (Fig 4A), indicative of homomeric histidine kinase interactions that reconstitute the split adenylate cyclase. Co-expression of the T25-SkaH fusion with either a T18c-LovK or T18c-SpdS* fusion also resulted in blue colonies (Fig 4A). B2H reporter strains co-expressing T25-LovK and T18c-SpdS* constructs yielded white colonies (Fig 4A). These data provide evidence that SkaH can directly interact with both LovK and SpdS to form a heteromeric kinase complex. However, LovK and SpdS do not directly interact in a B2H assay.

**Fig 4.**
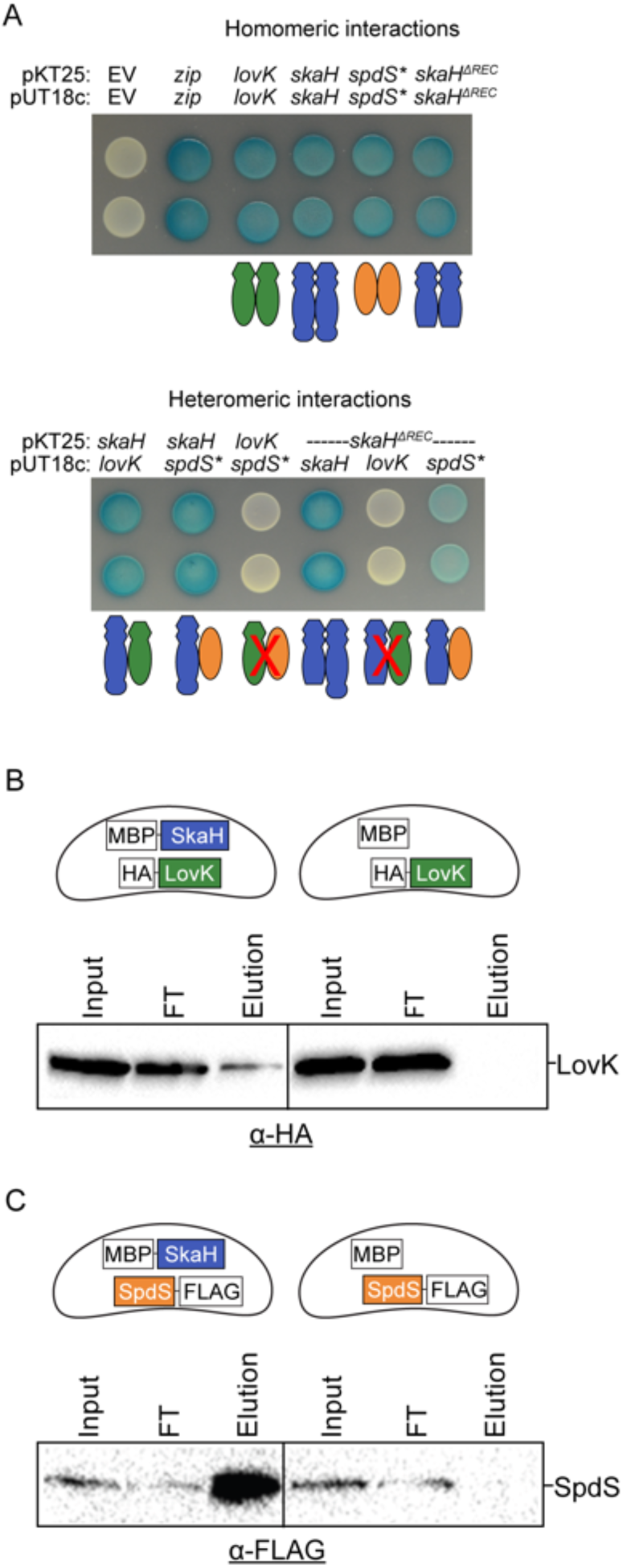
SkaH physically interacts with LovK and SpdS *in vitro* and *in vivo*. **A)** Bacterial two-hybrid (BTH) experiments to assess interactions between histidine kinase fusions to a split adenylate cyclase. Fusions with *spdS* lack the transmembrane domain, notated *spdS\**. *skaH*^Δ*REC*^ is a truncated version of SkaH that lacks the REC domain. Two biological replicates are shown for each co-expression combination. Protein-protein interaction of fusions reconstitutes the split adenylate cyclase encoded on pKT25 and pUT18c, and results in a blue color on agar plates containing x-gal. Strains expressing fusions that do not interact appear white. Zip = positive control. Histidine kinase protein-protein interaction models are schematized below each BTH assay as 1:1 interactions. However, the actual oligomeric state of detected interactions is not known. **B and C)** Co-purification experiments in which MBP is affinity purified with amylose resin and co-purifying kinases are detected by Western blot. The fusion proteins co-expressed in each experiment are schematized in the cells above each blot. Representative immunoblot against the HA and FLAG epitope tags are shown in B and C respectively. Input = clarified lysate, FT = Flow through.

SkaH is a hybrid histidine kinase, so we next tested whether the heteromeric kinase interactions observed in the B2H assay required the REC domain of SkaH. We fused a short version of SkaH that lacked the REC domain (SkaH^ΔREC^) to the T25 and T18c fragments of adenylate cyclase. Co-expression of SkaH^ΔREC^ fusions to both T25 and T18c resulted in blue colonies when grown on agar plates containing X-gal (Fig 4A); co-expression of T25-SkaH^ΔREC^ and T18c-SkaH also yielded blue colonies (Fig 4A) providing evidence that the homomeric interaction between SkaH monomers does not require the REC domain. A B2H reporter strain co-expressing T25-SkaH^ΔREC^ and T18c-LovK yielded white colonies (Fig 4A). Co-expression of T25-SkaH^ΔREC^ and T18c-SpdS* yielded light blue colonies, indicating the interaction was qualitatively weaker than the observed B2H interaction between full-length SkaH and SpdS* (Fig 4A). Together, these results provide evidence that the REC domain of SkaH is necessary for the heteromeric interaction with LovK observed in a B2H assay. The SkaH REC domain is not absolutely required for the B2H interaction we observe with SpdS*.

To test if the SkaH-LovK and SkaH-SpdS interactions observed in the heterologous B2H system occur *in vivo*, we tested whether LovK and/or SpdS co-purify with SkaH when affinity purified from *C. crescentus* lysate. Specifically, we ectopically expressed different combinations of histidine kinase fusions in *C. crescentus*: a maltose binding protein (MBP) N-terminally fused to SkaH (MBP-SkaH), a HA epitope tag N-terminally fused to LovK (HA-LovK), and a 3xFLAG tag C-terminally fused to full length SpdS (SpdS-3xFLAG). Upon co-expression in *C. crescentus*, MBP-SkaH was immobilized on an amylose affinity resin in an effort to pull down potential heteromeric kinase complexes. Co-purifying proteins were detected by Western blot. When MBP-SkaH was co-expressed with either HA-LovK or SpdS-3xFLAG, both fusions co-eluted with MBP-SkaH after a stringent wash and elution with maltose (Fig 4B and 4C). As a negative control, we co-expressed MBP alone with either HA-LovK or SpdS- 3xFLAG. Neither HA-LovK nor SpdS-3xFLAG co-eluted with MBP (Fig 4B and 4C), indicating that the interactions observed were not due to protein interaction with MBP. From these data, we conclude that SkaH physically interacts with both LovK and SpdS in *C. crescentus*.

### *spdR* regulates *hfiA* transcription indirectly

A major goal of our genetic selection was to identify direct regulators of *hfiA* transcription downstream of LovK. The DNA-binding response regulator, SpdR, emerged as a clear candidate from our data. We utilized genetic, molecular and genomic approaches to test the role of SpdR as a regulator of *hfiA* transcription. As described above, alanine substitution of the conserved phosphorylation site in the SpdR REC domain (D64A), phenocopied the Δ*spdR* deletion mutation in a *lovK*(H180A) background (Fig 3A). Moreover, a classic phosphomimetic substitution of aspartate to glutamate (D64E) in a wild-type background decreased activity from the P*_hfiA_-lacZ* transcriptional reporter and increased surface adhesion (Fig 5A). These data provide genetic evidence that SpdR can function to repress *hfiA* transcription and modulate cell surface adhesion in a phosphorylation dependent manner.

**Fig 5.**
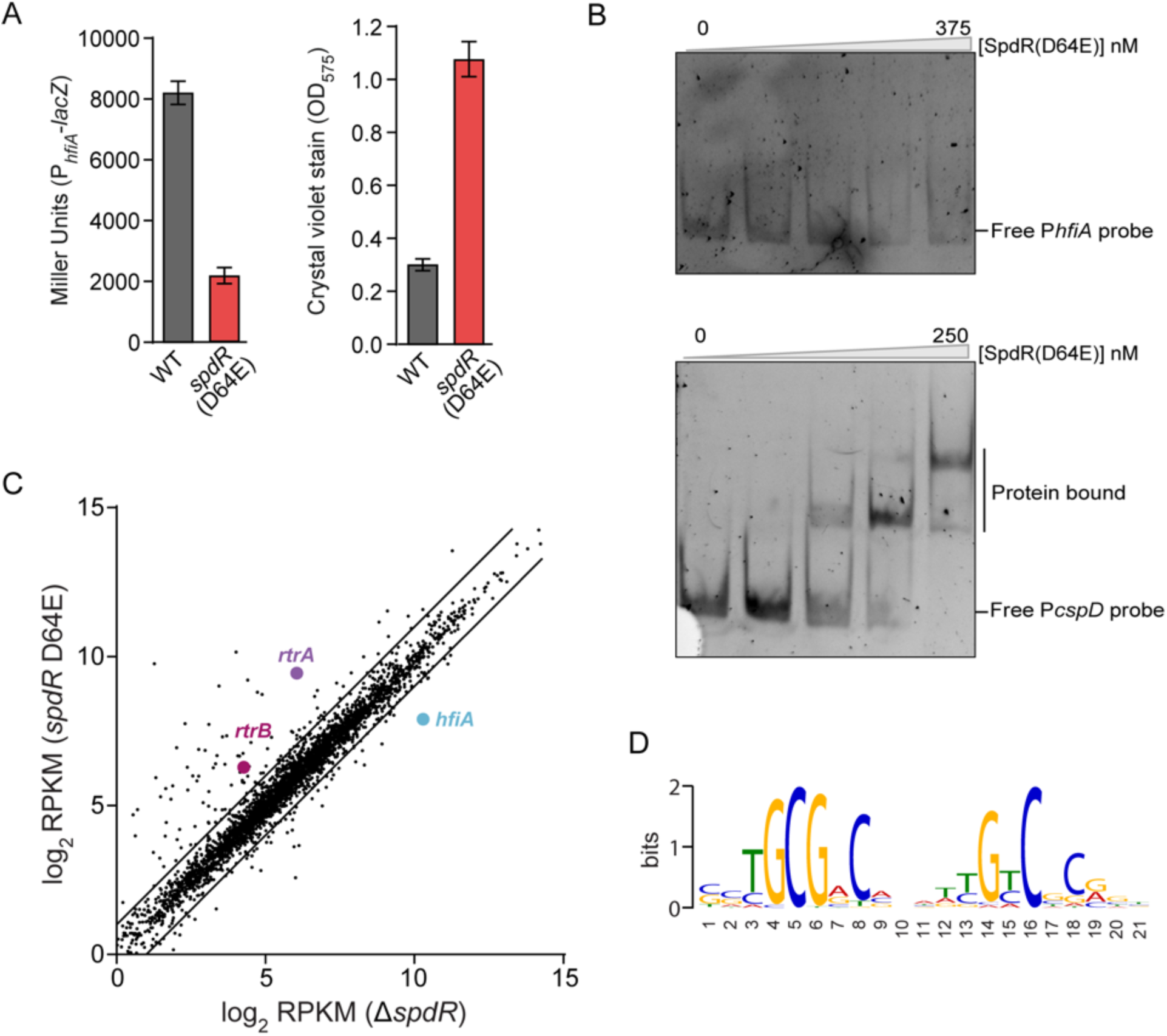
SpdR is an indirect repressor of hfiA trancription. **A)** β-galactosidase activity of P*_hfiA_-lacZ* transcriptional fusion (left), and attachment of cells to polystyrene measured by crystal violet stain (right) for wild-type (WT) and *spdR*(D64E) strains. Data represent mean ± s.d.; n=5-8. **B)** Electrophoretic mobility shift assay using recombinant His_6_-SpdR(D64E) and a fluorescently labeled hfiA promoter probe (top) or cspD promoter probe (bottom). His_6_-SpdR(D64E) protein concentrations when incubated with *hfiA* probe were 0, 50, 125, 250, 375 nM, and when incubated with *cspD* probe were 0, 25, 50, 125, 250 nM. Experiments were conducted three times. **C)** RNA-Seq analysis of transcript levels in spdR(D64E) and ΔspdR strain. Log_2_(mean RPKM) from each strain are plotted against each other where each gene is a dot. Genes of interest (*hfiA, rtrA, rtrB*) are marked. Lines indicate two-fold change cutoffs. See Methods for details. **D)** Promoter motif identified by MEME in genes positively regulated by *spdR*(D64E) compared to Δ*spdR*.

To test whether SpdR is a direct regulator of *hfiA*, we performed an electrophoretic mobility shift assay (EMSA) using a fluorescently labeled hfiA promoter probe and His_6_-SpdR(D64E) recombinant protein. Increasing concentrations of His_6_-SpdR(D64E) did not retard gel mobility of the *hfiA* probe (Fig 5B). As a positive control, we performed an EMSA using a fluorescently labeled probe corresponding to the promoter of *cspD*, a gene reported to be directly regulated by SpdR [39]. Migration of the labeled cspD promoter probe was retarded by increasing concentrations of His_6_-SpdR(D64E) and this probe displayed the same double shift as previously published (Fig 5B) [39]. These results provide evidence that SpdR does not bind the *hfiA* promoter, and therefore support a model in which SpdR indirectly regulates *hfiA* transcription.

### RNA sequencing analysis defines the SpdR regulon

Given the evidence that SpdR regulates *hfiA* transcription indirectly, we experimentally defined the transcriptional regulon of SpdR with the goal of uncovering direct *hfiA* regulators that are expressed downstream of SpdR. We measured steady-state transcript levels in a strain missing *spdR* (Δ*spdR*) and a strain expressing the active phosphomimetic *spdR*(D64E) allele by RNA sequencing (RNA-seq). The majority of genes with differential transcriptional profiles are more highly expressed in the constitutively active *spdR*(D64E) strain (Fig 5C and Table S2). Importantly, *hfiA* is among the small set of transcripts that are downregulated in *spdR*(D64E) relative to Δ*spdR* (5.1-fold, p < e^−16^) (Fig 5C and Table S2). A consensus binding motif for SpdR has been defined in *C. crescentus* [39, 40] and for SpdR homologs in other species [46–48]. To predict direct targets of SpdR, we utilized the MEME motif discovery platform [49] to identify shared sequence motifs in the promoter regions (−150 to +50) of all the genes with at least 2-fold expression differences between *spdR*(D64E) and Δ*spdR*. This approach revealed a predicted SpdR binding site (Fig 5D) in the promoter regions of 82 genes (Table S2), suggesting that these genes are directly regulated by SpdR. A scan of the region from −500 bp to +100 bp around the start *hfiA* codon failed to identify a SpdR binding sequence. These data provide additional evidence that SpdR is not a direct repressor of *hfiA* transcription.

### An approach to identify direct *hfiA* transcriptional regulators downstream of SpdR

We hypothesized that transcription factor(s) regulated downstream of SpdR were responsible for repression of *hfiA* transcription. We took a candidate approach and examined all the differentially expressed genes (>2-fold) in our RNA-Seq dataset that contained a predicted DNA-binding domain. Each of the six predicted transcription factors in this set were activated by SpdR (Table S2). Thus, we predicted that the repressive effect of SpdR on *hfiA* transcription was the result of upregulation of one or more transcription factors that directly repress *hfiA* (Fig 6A). We individually overexpressed these transcription factors and measured *hfiA* promoter activity and surface attachment (not shown). Two *<u>X</u>*enobiotic *<u>R</u>*esponse *<u>E</u>*lement (XRE)-family transcription factors, *CC_3164* (*CCNA_03267*) and *CC_2330* (*CCNA_02415*), displayed the overexpression phenotypes expected for an *hfiA* repressor: a decrease in transcription from the *hfiA* promoter (Fig 6B) and an increase in surface adhesion (Fig S8), compared to wild-type. *CC_3164* was a more potent *hfiA* repressor than *CC_2330*. Coordinate overexpression of *CC_3164* and *CC_2330* did not decrease *hfiA* expression more than *lovK*(H180A) or a strain overexpressing only *CC_3164* (Fig 6B). Strains overexpressing either one or both *CC_3164* and *CC_2330* had the same surface adhesion phenotype as *lovK*(H180A) (Fig S8). Importantly, the promoter regions of *CC_3164* and *CC_2330* each contain a predicted SpdR binding site (Table S2), providing evidence that each are directly regulated by SpdR. Since *CC_3164* and *CC_2330* are in the SpdR regulon and regulate *hfiA* promoter activity, we named these genes *rtrA* and *rtrB,* for *<u>R</u>*eg*BA t*ranscriptional *r*egulator *<u>A</u>* and *<u>B</u>*, respectively.

**Fig 6.**
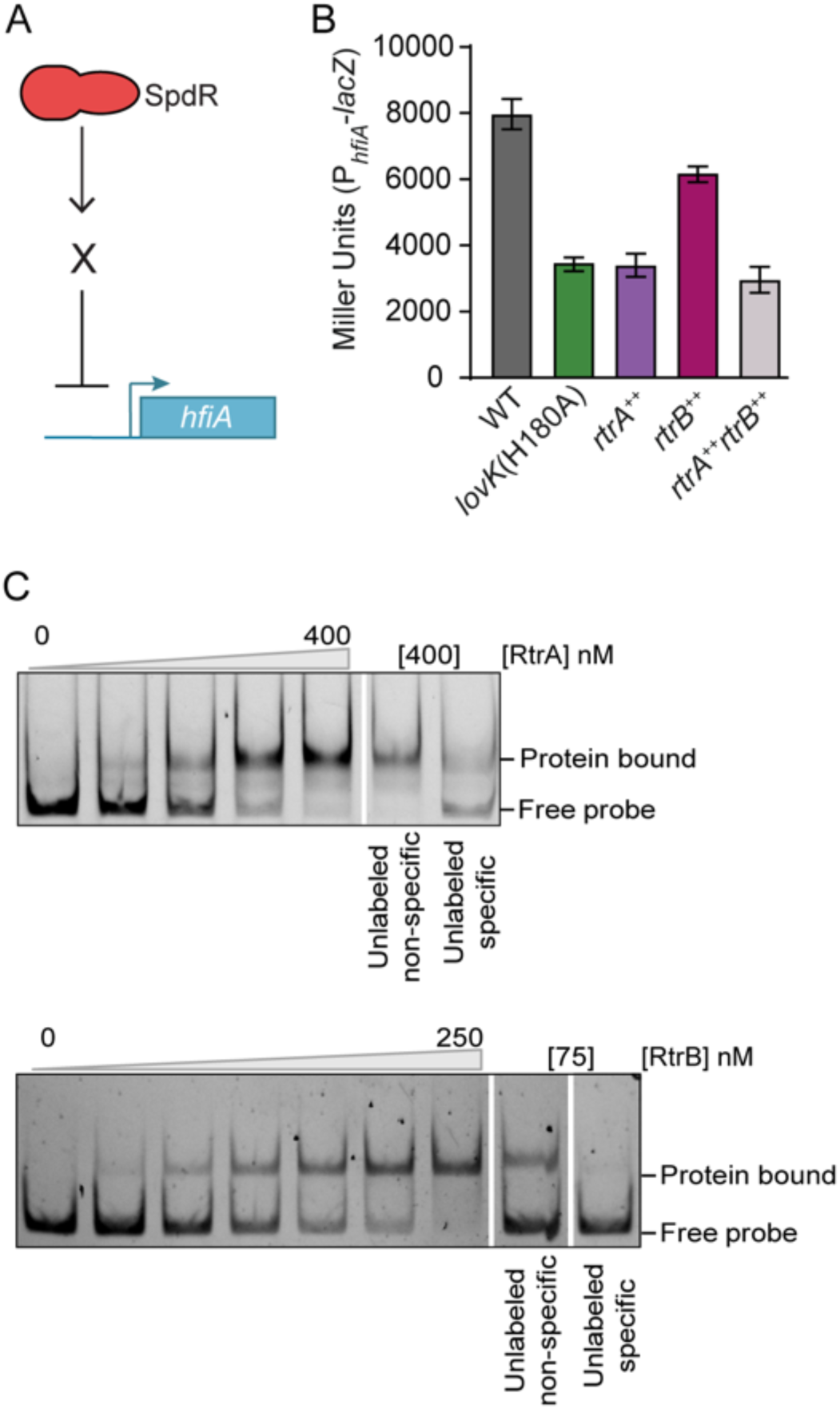
*rtrA* (*CC_3164*) and *rtrB* (*CC_2330*) are direct repressors of *hfiA*. **A)** We postulated that transcription factor(s) X, downstream of SpdR, repressed *hfiA* transcription. **B) β**-galactosidase activity from P*_hfiA_-lacZ* transcriptional reporter in different genetic backgrounds. Wild-type (WT) and *lovK*(H180A) carry empty vectors pMT680 and pMT585 integrated at the xylose locus as controls. *rtrA*^++^ carries pMT680-*rtrA* and pMT585 empty vector. *rtrB*^++^carries pMT585-*rtrB* and pMT680 empty vector. *rtrA*^++^*rtrB*^++^ carries pMT680-*rtrA* and pMT585-*rtrB*. Genes in both plasmids are expressed from a xylose inducible promoter. Data represent mean ± s.d.; n=6. **C)** Electrophoretic mobility shift assay of purified RtrA (top) and His_6_-RtrB (bottom) with a fluorescently labeled hfiA promoter probe, with increasing concentrations of protein (0, 100, 200, 300, 400 nM for RtrA and 0, 10, 25, 50, 75, 100, 250 nM for His_6_-RtrB). Unlabeled specific and non-specific probes are in 10-fold excess of the labeled hfiA promoter probe. Blots are representative of at least three independent experiments.

**Fig 7.**
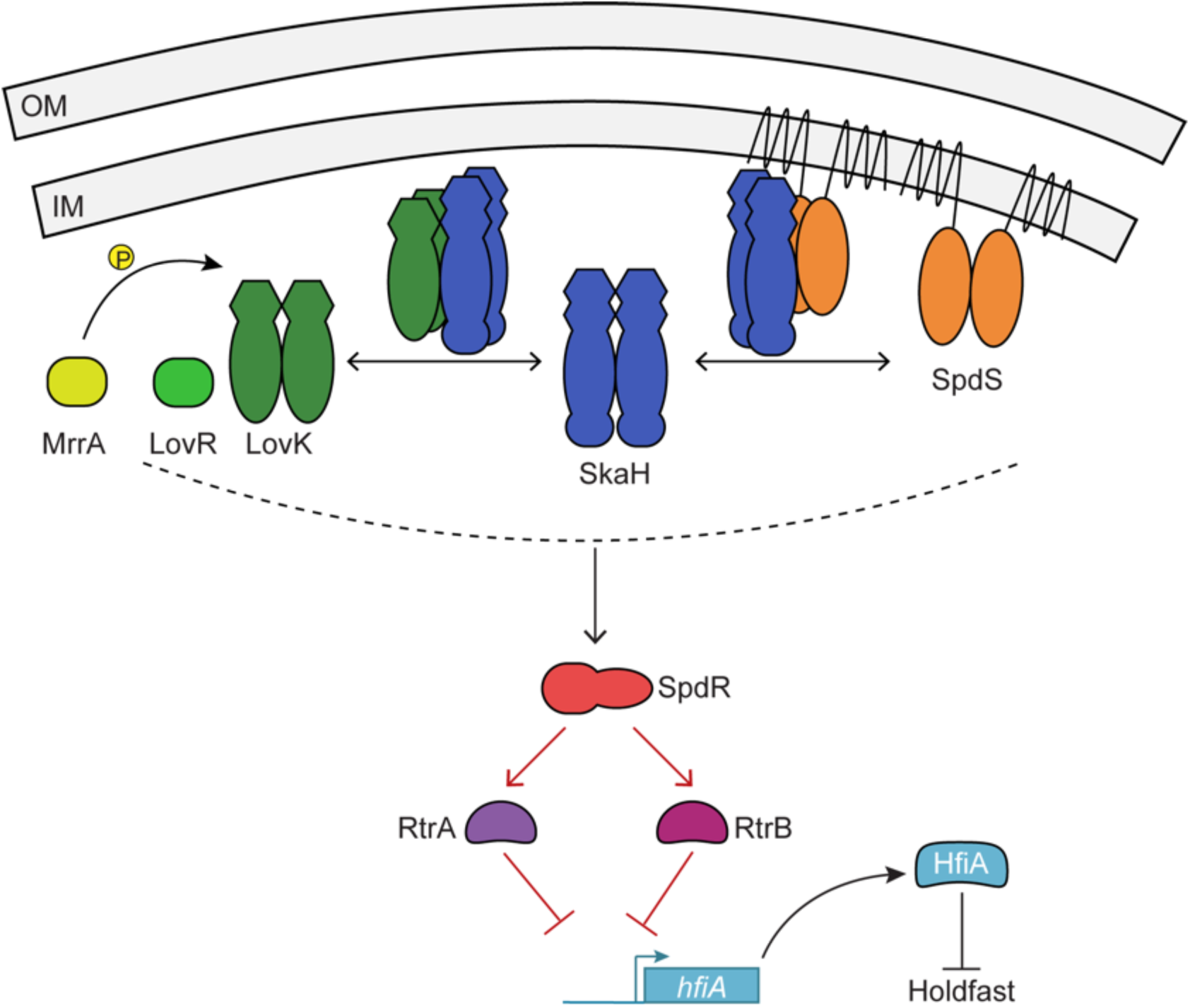
Network model of TCS proteins and transcription factors that regulate *hfiA* transcription and surface adhesion. Proposed model in which the hybrid histidine kinase, SkaH, interacts with both LovK and SpdS to mediate *hfiA* transcription. The oligomeric state of these kinase complexes is not known. Interactions and/or phosphoryl transfer between the kinases and response regulators are proposed to collectively regulate SpdR phosphorylation. Phosphorylated SpdR activates transcription of RtrA and RtrB, transcription factors that directly bind to and repress the *hfiA* promoter. The SDRR, MrrA, can transfer a phosphoryl group to LovK and also modulates *hfiA* transcription in a LovK dependent manner [16]. Red lines indicate direct transcriptional regulation. Dashed line indicates the unknown flow of phosphoryl groups.

### RtrA and RtrB are direct regulators of *hfiA* transcription

We next tested whether RtrA and RtrB directly bind the *hfiA* promoter by EMSA. Increasing concentrations of either protein retarded mobility of a labeled *hfiA* promoter probe relative to the probe alone (Fig 6C). To assess the specificity of this interaction, we added 10-fold excess of unlabeled specific probe or unlabeled non-specific probe. The unlabeled specific probe competed with the labeled probe for binding to RtrA and RtrB, while unlabeled non-specific probe had little effect on binding of RtrA or RtrB to the *hfiA* promoter region (Fig 6C). These data support a model in which RtrA and RtrB directly repress *hfiA* transcription.

## Discussion

Adherence of a *C. crescentus* cell to a surface via its holdfast is effectively permanent. As such, it is not surprising that holdfast synthesis is a highly regulated process. *hfiA* is a potent inhibitor of holdfast synthesis, and its transcription is controlled by a suite of cell cycle and environmental regulatory proteins [14–16]. In this study, we sought to understand the molecular mechanism by which the TCS, LovK-LovR, controls *hfiA* transcription. A forward genetic selection followed by candidate gene studies led to the discovery of a network of TCS proteins and transcription factors that coordinately function with LovK to regulate *hfiA* transcription. Specifically, we have shown that the hybrid histidine kinase, SkaH, forms heteromeric complexes with the LovK kinase and the SpdS kinase *in vivo*. This suite of kinases works through the DNA-binding response regulator, SpdR, to activate expression of two transcription factors, RtrA and RtrB, which directly bind the *hfiA* promoter and repress its transcription. Our data provide evidence for a multi-component sensory system that has the capacity to detect, and perhaps integrate, multiple signals to regulate holdfast development.

### Signaling through the LovK-LovR system

*C. crescentus lovK-lovR* overexpression modulates cell adhesion by influencing *hfiA* transcription [14], but it also indirectly represses dozens of genes in the General Stress Response (GSR) regulon [50] through an independent molecular mechanism [14]. The mechanism by which the SDRR LovR functions with LovK to regulate *hfiA* and the GSR remains unclear, but previous genetic data have suggested that its function may be to simply dephosphorylate LovK [33, 50]. This model is based on the result that overexpression of either *lovK* or *lovR* alone has no effect on cell-to-cell adhesion [33] or on repression of GSR transcription [50], while coordinate overexpression represses GSR transcription [50], enhances cell-to-cell [33] and surface adhesion [14], and represses *hfiA* transcription [14]. These data support a model in which LovK is active in its unphosphorylated state. Similarly, our analysis of *hfiA* transcription and surface attachment in a strain that solely expresses the non-phosphorylatable *lovK*(H180A) allele supports such a model (Fig S1).

What, then, occurs at the molecular level that explains these phenotypes? Unphosphorylated-LovK may function as an allosteric regulator of other kinases or kinase complexes that control SpdR phosphorylation. LovK may also dephosphorylate a receiver protein or receiver domain required for SpdR phosphorylation. There is evidence that LovK can function as a dephosphorylase in the case of the SDRR MrrA, which was recently proposed to serve as a signaling hub that controls GSR transcription. MrrA also enhances *hfiA* expression in a manner that requires *lovK*, but not *lovR* [16]. These data suggest MrrA is an additional TCS protein that functions upstream of LovK in the network of TCS proteins that modulate *hfiA* transcription. We hypothesize that non-phosphorylatable LovK(H180A) leads to high levels of phosphorylated SpdR, which results in repression of *hfiA* transcription. Given the requirement for SkaH, we do not favor a model in which LovK directly phosphorylates or dephosphorylates SpdR. One possibility is that LovK directly dephosphporylates the SkaH REC domain; through an unknown mechanism, perhaps involving interactions with SpdS, this would lead to phosphorylated SpdR. This model is supported by the requirement of the conserved aspartate of SkaH(D550) for the lovK(H180A) adhesion and P*_hfiA_-lacZ* transcription phenotypes, and the requirement of the REC domain for HK heteromeric interactions by B2H (Fig 3 and 4).

### Interplay between HisKA and HWE/HisKA_2 families of histidine kinases

SkaH and LovK belong to the HWE/HisKA_2 histidine kinase family, which is defined by unique primary structure features in the dimerization, histidine phosphotransfer (DHp), and catalytic (CA) domains [35, 36]. HWE/HisKA_2 proteins are reported to function as part of complex signal transduction systems, including the GSR in *Alphaproteobacteria* [16, 26] and cyst development in *Rhodospirillum centenum* [51, 52], and have been reported in unusual oligomeric forms including monomer [53] and hexamer [54]. Though SpdS is a member of the more common HisKA family of HKs, its homolog in *Rhodobacter capsulatus* (RegB) can form tetramers mediated by an intermolecular disulfide bond that forms through a conserved cysteine [55]. It is not known if *C. crescentus* SpdS forms tetramers, but we predict that it may since it contains the conserved cysteine (C248) present in *R. capsulatus* RegB. It is therefore possible that HWE/HisKA_2 kinases (SkaH and LovK) and a HisKA kinase (SpdS) form unusual, high-order oligomers in the cell, and that changes in the oligomeric states of these proteins affect their function as regulators of *hfiA* and GSR transcription. Though we have presented clear evidence for interactions between SkaH, LovK, and SpdS, the structural basis of these interactions remains undefined. It is important to consider the possibility that the heteromeric HK interactions we observe under our assay conditions do not reflect the full extent of kinase interactions in the cell. Specifically, it is possible that the LovK-SkaH-SpdS complex is part of an even higher order complex of regulatory proteins, as the case for the ≈2 MDa stressosome complex that controls the σ^B^-dependent general stress response (GSR) in *Bacillus subtilis* [56, 57].

Finally, our data provide evidence for an emerging model in which HisKA and HWE/HisKA_2-family HKs can function together to regulate gene expression. A previous study showed that the SDRR, MrrA, can be phosphorylated and/or dephosphorylated by kinases from both of these families of HKs in *C. crescentus* [16]. Furthermore, deletion of all the HWE/HisKA_2 HKs in *Sphingomonas melonis* Fr1 does not completely ablate GSR transcription [26], suggesting that one or more HisKA kinases also play a role in regulating this system. This study provides additional evidence for complex signal transduction systems involving both HisKA and HWE/HisKA_2 histidine kinase families.

### A consortium of sensors regulate *hfiA* to control cell adhesion

Crystal violet staining data reveal a moderate but reproducible defect in surface adhesion when *skaH* or *spdR* are deleted in a wild-type background (Fig 3D and Fig S5). This defect is more pronounced as culture density increases (Fig 3D). We note that small differences in surface adhesion are challenging to measure. Nevertheless, our results are consistent with a recent genome-scale analysis of adhesion to a cellulosic surface where mutants with disruptions in *skaH*, *spdS*, *spdR*, and *rtrB* exhibited reduced cell attachment to cheesecloth in complex medium (Fig S9) [58]. This independent genetic experiment supports a model in which these gene products function together as positive regulators of surface attachment across distinct media conditions.

The specific environmental or intracellular signals that regulate activity of the sensory systems identified in our selection are not known. However, SpdS-SpdR homologs have been studied extensively and shown to sense the cellular redox state by three mechanisms: *1)* binding of oxidized quinone [59, 60], *2)* cysteine sulfenic acid modification [61] or *3)* disulfide bond formation [55] at a conserved reactive cysteine. *C. crescentus* SpdS transfers a phosphoryl group to SpdR *in vitro* [39], but the role of SpdS-SpdR phosphorylation *in vivo* remains unclear. In other *Alphaproteobacteria*, SpdS homologs have been reported to dephosphorylate their cognate regulators [62–65], and thus we cannot exclude the possibility that SpdS may primarily function to dephosphorylate SpdR in *C. crescentus* cells. In fact, we observed that upon entering stationary phase, a Δ*spdS* mutant has dramatically enhanced surface adhesion, similar to the hyper-adhesive phosphomimetic *spdR*(D64E) mutant. This stationary phase Δ*spdS* phenotype suggests a model in which stationary phase signal(s) activate SpdS as a SpdR~P dephosphorylase. This model remains to be tested.

*C. crescentus* LovK has the capacity to function as both a redox sensor [37] and a photosensor [33]. The genetic interactions we uncovered between LovK and SpdS-SpdR may reflect a fundamental role for LovK in redox sensing *in vivo*. The ability of SkaH to function as a sensor has not been characterized, though a database search using the Phyre2 profile-profile alignment algorithm [66] provides evidence for two PAS-like sensor domains [67] N-terminal to the HWE kinase domain. The multi-protein regulatory system we report here could potentially sense a variety of environmental signals (e.g. redox potential, light, small molecules) to control *hfiA* expression.

### Downstream regulators that directly control *hfiA* transcription

We identified two transcription factors, *rtrA* and *rtrB*, that function downstream of SpdR to directly repress *hfiA* transcription and control surface attachment. *rtrA* was previously identified as part of the stationary phase SpdR regulon in *C. crescentus* [40], while *rtrB* was not. Our transcriptomic experiments were conducted on cells in logarithmic phase expressing a phosphomimetic SpdR allele. Studies of RegA-dependent transcription in *Rhodobacter capsulatus* have shown that the composition of the RegA regulon depends on the growth medium [68]. Similar conditional regulation by SpdR may exist in *C. crescentus*, and thus *rtrA* and *rtrB* may be activated and control *hfiA* transcription under distinct sets of environmental conditions. Efforts to globally define the regulons of *rtrA* and *rtrB* are ongoing and will provide a more complete understanding of the broader functional roles of these *hfiA* regulators in *C. crescentus*.

## Methods

### Strain growth conditions

*Escherichia coli* strains were cultivated in Lysogeny Broth (LB) or LB agar plates (1.5% agar) at 37°C. Antibiotics were added to the following final concentrations as required: kanamycin 50 µg/ml, chloramphenicol 20 µg/ml, ampicillin 100 ug/ml, tetracycline 12 µg/ml, gentamicin 15 µg/ml and spectinomycin/streptomycin 50 µg/ml / 30 µg/ml.

*Caulobacter crescentus* strains were cultivated on peptone-yeast extract (PYE) agar (0.2% peptone, 0.1% yeast extract, 1 mM MgSO_4_, 0.5 mM CaCl_2_, 1.5% agar) at 30°C [69]. Liquid cultures of C. crescentus were cultivated at 30°C in PYE (0.2% peptone, 0.1% yeast extract, 1 mM MgSO_4_, 0.5 mM CaCl_2_) or M2 defined medium supplemented with xylose (0.15%) as the carbon source (M2X) [69]. For expression of a gene from the *vanA* promoter, a final concentration of 0.5 mM of vanillate was added to M2X (M2VX). Antibiotics were added to the following final concentrations as required for liquid cultures: tetracycline 1 µg/ml. For solid medium growth, antibiotics were added as follows: kanamycin 25 µg/ml, chloramphenicol 1 µg/ml, tetracycline 2 µg/ml, gentamicin 5 µg/ml and spectinomycin/streptomycin 50 µg/ml / 5 µg/ml.

### Plasmid construction

*C. crescentus* DNA was amplified from colonies using KOD Xtreme hot-start polymerase (EMD Biosciences/Novagen). PCR reactions were supplemented with 5% dimethyl sulfoxide (DMSO). All plasmids were cloned in *E. coli* Top10 (Invitrogen, Carlsbad, CA). The sequences of all cloned products were confirmed in the target plasmids.

Gene deletions and point mutations were generated by spliced overlapping PCR using external primers with specific restriction enzyme sites. PCR products were digested and ligated into the pNPTS138 plasmid [70] at the necessary restriction enzyme sites. Plasmids for ectopic gene expression were generated by ligating a digested PCR product of the gene of interest into *colE1*-based integrating plasmids with either a vanillate (P*_van_*) or xylose (P*_xyl_*) inducible promoter [71]. Plasmids are listed in Table S3.

### Strain construction

Chromosomal allele replacements were made using standard two-step recombination using *sacB* for counterselection. Briefly, pNPTS138-derived plasmids were transformed into *C. crescentus* by electroporation. Primary integrants were selected by growth on PYE Kan plates, inoculated into liquid PYE and grown overnight without selection. Cultures were plated on PYE agar supplemented with 3% sucrose to select for clones that lost the plasmid in a second recombination event. Chromosomal allele replacements were confirmed by PCR amplification of the gene of interest from kanamycin sensitive clones. pRKlac290 transcriptional reporter plasmid P*_hfiA_-lacZ* was conjugated into *C. crescentus* by triparental mating [69]. *C. crescentus* was then grown on PYE plates supplemented with tetracycline to select for cells carrying the plasmid and nalidixic acid to counterselect against *E. coli*. Integrating expression plasmids were transformed into *C. crescentus* by electroporation. Strains carrying the plasmid were selected on PYE agar supplemented with a plasmid-specific antibiotic.

### Protein purification for EMSA

For heterologous expression, pET-based plasmids were transformed into Rosetta(DE3)pLysS (Novagen). Strains were inoculated into 10 ml of LB liquid medium supplemented with Kan 50 µg/ml and grown overnight shaking at 37°C. The overnight culture was used to inoculate a flask with 250 ml of fresh LB liquid medium supplemented with Kan 50 µg/ml and grown shaking to OD_600nm_ of 0.6 - 0.8 at 37°C. Then, a final concentration of 1 mM isopropyl **β**-D-1-thiogalactopyranoside (GoldBio) was used to induce protein expression for 4 hrs under the same conditions. The cultures were harvested by centrifugation at 11,000 × g for 7 min at 4°C, resuspended in 15 ml of resuspension buffer (10 mM Imidazole, 125 mM NaCl, 25 mM Tris pH 7.6) and saved at −20°C until needed. When samples were thawed, DNAseI and PMSF were added to final concentrations of 250 µg/ml and 800 µM, respectively. The cells were then disrupted by one passage in a microfluidizer (Microfluidics LV1) and the resulting lysate was clarified by centrifugation at 39,000 × g for 15 min at 4°C. Purification of His_6_-RtrB, His_6_-SUMO-RtrA and His_6_-SpdR(D64E) was performed by nickel affinity chromatography (nitrilotriacetic acid resin; GE Healthcare). After binding of the clarified lysate to the column, the sample was washed with 30-50 ml of resuspension buffer followed by 10 ml of wash buffer (75 mM Imidazole, 125 mM NaCl, 25 mM Tris pH 7.6). Protein was eluted with 1-2 ml of elution buffer (500 mM Imidazole, 125 mM NaCl, 25 mM Tris pH 7.6). Sample purity was assessed by resolving the purified protein by 12% SDS-PAGE and the gels were stained with Coomassie blue.

For purification of RtrA, ULP1 protease was added to the eluted protein to cleave the His_6_SUMO tag and this sample was dialyzed overnight at 4°C in 1 L of buffer containing 25 mM Tris pH 7.6, 125 mM NaCl_2_, 50% glycerol. The cleaved protein was further purified by nickel affinity chromatography, where the flow through containing the cleaved untagged RtrA was collected for use in this study. For His_6_-RtrB purification, the eluted protein was dialyzed overnight at 4°C in 1 L of buffer containing 25 mM Tris pH 7.6, 200 mM KCl, 10 mM MgCl_2_, 15mM EDTA pH 8.0 and 50% glycerol. For purification of His_6_-SpdR(D64E), the eluted protein was dialyzed overnight at 4°C in 1 L of buffer containing 25 mM Tris pH 7.6, 125 mM NaCl_2_ and 50% glycerol. Aliquots of the purified proteins were saved at −80°C.

### Genetic selection

The goal of this selection was to isolate spontaneous mutants in which the *lovK*(H180A) allele could no longer repress the hfiA promoter (P*_hfiA_*). We used a lovK(H180A) strain carrying an integrated plasmid in which the promoter of *hfiA* was fused to cat (P*_hfiA_-cat*). The cat gene encodes chloramphenicol acetyltransferase, which confers resistance to the antibiotic chloramphenicol in a dose dependent manner. We used growth rate in the presence of chloramphenicol as a proxy for P*_hfiA_* activity and enriched for spontaneous mutants that grew faster than the parental *lovK*(H180A) strain in the presence of chloramphenicol. We identified optimal selection conditions by comparing growth rate of wild-type and lovK(H180A) strains carrying the P*_hfiA_-cat* plasmid in a range of chloramphenicol concentrations. The highest dynamic range between growth rates in the wild-type and *lovK*(H180A) backgrounds was at 1 - 2 µg/ml chloramphenicol (data not shown).

For each independent selection, one ml of an overnight culture of the parental strain was used to inoculate flasks containing 500 ml of M2X medium supplemented with a final concentration of 1 - 2 µg/ml of chloramphenicol. Flasks were incubated shaking at 30°C until they reached OD_660nm_ of 0.6 - 0.9. At this point, 100 - 500 µl of the cultures was used to inoculate 500 ml of fresh M2X medium. Cultures were allowed to grow and this passaging process was performed until the diluted cultures grew fast enough to reach OD_660nm_ of 0.6 - 0.9 in a single overnight incubation. Upon initial inoculation, cultures took 3 – 5 days to reach the desired OD_660nm_ but after 5 - 7 serial passages cultures only took about 18 hrs to reach OD_660nm_ of 0.6 - 0.9. This indicated that the population was more resistant to chloramphenicol. Individual mutant clones were isolated from the final (fast-growing) culture by plating serial dilutions on PYE agar supplemented with gentamicin 5 µg/ml, which selected for cells carrying the P*_hfiA_-cat* plasmid. Several isolated single colonies were picked for further analysis.

Several secondary screens were performed on the isolated mutants. First, we sequenced the hfiA promoter from the P*_hfiA_-cat* plasmid in order to distinguish clones with intact selection systems from those with *cis* mutations in the *hfiA* promoter. We then confirmed faster growth rate in the presence of chloramphenicol. Then, to distinguish mutants that result in de-repression of P*_hfiA_* from those that confer chloramphenicol resistance through other mechanisms, we evaluated bulk surface attachment of the isolated mutants by crystal violet stain or ring assay (see below) as a proxy for activity of the native *hfiA* promoter. Finally, mutants that displayed faster growth rate and decreased surface attachment compared to the lovK(H180A) parental strain were transformed with the P*_hfiA_-lacZ* transcriptional fusion plasmid. **β**-galactosidase activity was measured to confirm de-repression of the *hfiA* promoter in the spontaneous mutants compared to the *lovK*(H180A) strain. Based on the results obtained during the secondary screens, several isolates were selected and the mutations in these strains were mapped by whole genome sequencing.

### Whole genome sequencing

We isolated genomic DNA from the *C. crescentus* parent and the mutant strains arising from our chloramphenicol selection using a standard guanidinium thiocyanate extraction and isopropanol/ethanol precipitation. The DNA was randomly sheared and libraries were prepared for whole genome shotgun sequencing using an Illumina HighSeq 2500 (50-bp single end reads). Whole-genome sequence data from the parent and mutant strains were assembled to the *C. crescentus* NA1000 genome (genbank accession CP001340), and polymorphisms between the parental strain and suppressor strains were identified using Geneious v 11 [72], see Table S1. The sequenced genome libraries yielded an average of 11.4 million reads, resulting in global average depth of coverage of 141x.

### Crystal violet stain surface attachment assay

Colonies from PYE agar plates were inoculated into 2 ml of M2X broth and grown overnight at 30°C shaking at 200 rpm. Overnight liquid cultures were diluted to a final OD_660nm_ of 0.00625 into wells of 24-well polystyrene plates containing 1 ml of M2X or M2VX. Plates were grown shaking at 155rpm at 30°C for varying times periods as indicated in the results. Plates were removed from the incubator and surface attached cells were measured with crystal violet stain. Briefly, the cultures were discarded and the wells were thoroughly washed with water. Then, 1.5 ml of 0.01% crystal violet in water was added to each well and the plates were incubated with shaking at 155 rpm for 5 min. The crystal violet solution was discarded and the wells thoroughly washed with water again. To dissolve the stain, 1.5 ml of 100% EtOH was added to each well and the plates incubated again with shaking for 5 min. Crystal violet stain extracted in each well was quantified by measuring absorbance at 575nm.

### B-galactosidase assay

Strains were inoculated in M2X medium from colonies on PYE-agar plates and grown shaking overnight at 30°C. Overnight cultures were then diluted back into fresh M2X to an OD660nm of 0.00025 – 0.00075 and grown in a shaking incubator at 30°C until they reached OD_660nm_ 0.05 - 0.15 (15 – 20 hrs). At this point, **β**-galactosidase activity was measured as previously described [14].

### Bacterial two-hybrid

We utilized a system previously described [42]. We co-transformed plasmids bearing fusions to either the T18c or T25 domains of adenylate cyclase into the adenylate cyclase null strain, BTH101, by electroporation. An aliquot of the outgrowth was plated on LB agar supplemented with Amp 100 µg/ml, Kan 50 µg/ml, X-gal 80 µg/ml and IPTG 0.5 mM and grown at 30°C for 24 hrs. Strains were inoculated into LB liquid medium supplemented with Amp 100 µg/ml, Kan 50 µg/ml and IPTG 0.5 mM and grown shaking overnight at 30°C. Overnight cultures were diluted to OD_600nm_ of 0.05 and 5 µl of each culture were spotted into LB agar plates supplemented with Amp 100 µg/ml, Kan 50 µg/ml, X-gal 80 µg/ml and IPTG 0.5 mM. The color of each spot was evaluated after 36 hours of growth at 30°C.

### Protein pulldown from cell lysate

*C. crescentus* colonies from PYE agar plates were inoculated into 2 ml of M2X and grown shaking overnight at 30°C. These cultures were diluted into 40 ml of M2X and grown until they reached OD_660nm_ of 0.4 - 0.6. The cultures were then induced with 0.5 mM vanillate for 3 hrs. The cells were harvested by centrifugation at 11,000 × g for 15 min at 4°C, resuspended in 6 ml of standard buffer (25 mM Tris pH 7.6, 125 mM NaCl) and frozen at −20°C. When thawed, cells were disrupted by one passage in a microfluidizer (Microfluidics LV1), PMSF was added to a final concentration of 800 uM and the lysates were cleared by centrifugation at 21,000 × g for 5 min at 4°C. The clear lysate was loaded into a column of approximately 200 µl amylose resin (NEB) prequilibrated in buffer, and allowed to bind. The resin was washed with 30-50 ml standard buffer and eluted with 150 µl standard buffer supplemented with 40 mM maltose. All the fractions collected for analysis were mixed 1:1 with SDS loading buffer, boiled for 2 min at 95°C and saved at −20°C for further Western Blot analysis.

### Western blot of pulldown fractions

15 μl of each pulldown fraction sample was loaded onto a Mini-PROTEAN TGX Precast 4-20% Gradient Gel (Bio-Rad). Samples were resolved at 35 mA constant current in SDS running buffer (0.3% Tris, 18.8% Glycine, 0.1% SDS). Proteins in the gel were transferred to an Immobilon®-P PVDF Membrane using a Mini Trans-Blot® Cell after preincubation in Western transfer buffer (0.3% Tris, 18.8% Glycine, 20% methanol). Transfer was carried out at 100 volts for 1 hr at 4°C in Western transfer buffer. The membrane was then blocked in 5% powdered milk in Tris-buffered Saline Tween (TBST: 137 mM NaCl, 2.3 mM KCl, 20 mM Tris pH 7.4, 0.1% Tween 20) shaking for 1 hr at 4°C. Incubation with the primary antibody, FLAG monoclonal antibody (clone FG4R) or HA-Tag monoclonal antibody, was carried out shaking overnight in 5% powdered milk TBST at 4°C. Membrane was then washed 3 times in TBST for a total of 30 min at room temperature. Incubation with Goat anti-Mouse IgG (H+L)-HRP secondary antibody was at room temperature for 1 h in TBST. Finally, the membrane was washed 3 times in TBST for a total of 30 min at room temperature. Chemiluminescence was performed using the SuperSignal™ West Femto Maximum Sensitivity Substrate (Pierce) and was imaged using a ChemiDoc MP imaging system (Bio-Rad). Chemiluminescence was measured using the ChemSens program with an exposure time of 30 - 60 sec.

### Electrophoretic Mobility Shift Assays

Promoter regions of interest were amplified by PCR using a set of appropriate primers in which the reverse primer was fluorescently labeled with AlexaFluor 488 (Integrated DNA Technology, DTT). The *hfiA* promoter probe was 275 bp (primers F:TGGTGGTCCTGATCCTCCTG and R: CACTGACAACATCCTGTCCG) and the *cspD* promoter probe was 199 bp (primers F: CTAGGGACTGCCATCTTCGG and R: TCGTAACCAGACATCCCACC). Specific cold specific competitor probe represents the same region as the corresponding labeled probe amplified with unlabeled primers with the same sequence. The cold non-specific competitor was a 220 bp PCR product amplified from the pNPTS138 plasmid using primers F: GTAAAACGACGGCCAG and R: CAGGAAACAGCTATGAC.

For reactions in which the binding of His_6_-SpdR(D64E) was tested, DNA binding reactions were performed in 20 µl reaction volumes containing binding buffer (20mM HEPES pH 7.8, 4 mM MgCl_2_, 37 mM KCl, 1 mM CaCl_2_, 0.75 mM dithiothreitol, 8% glycerol and 75 µg/ml bovine serum albumin), 8 ng of fluorescently labeled DNA probe and increasing concentrations of His_6_-SpdR(D64E) (0, 50, 125, 250, 375 nM for incubation with the *hfiA* promoter probe and 0, 25, 50, 125, 250 nM for incubation with the *cspD* promoter probe). Reactions were incubated at room temperature for 30 min in the dark. Then, 12.5 µl of each reaction was loaded on a fresh 6% native acrylamide gel pre-run for at least 30 min in 1X Tris Acetate EDTA buffer (TAE: 40 mM Tris, 20 mM acetic acid and 1 mM EDTA) in the dark at 4°C for 1 hr at 70 volts followed by 1.5 hr at 90 volts. The gels were imaged using the BioRad Chemidoc MP imaging system with a 60 - 120 sec exposure and the manufacturer’s settings for AlexaFluor 488 detection.

For reactions where the binding of RtrA was tested, DNA binding reactions were performed in 20 µl reaction volumes containing binding buffer (10 mM Tris pH7.6, 2 mM MgCl_2_, 50 mM KCl, 2 mM CaCl_2_, 0.2 mM dithiothreitol, 5% glycerol, 1 mM EDTA pH 8.0 and 1 mg/ml bovine serum albumin), 8 ng of fluorescently labeled DNA probe and increasing concentrations of RtrA (0, 100, 200, 300, 400 nM). For competition experiments, a 10-fold excess of cold competitor probe was added to the reaction. Reactions were incubated at room temperature for 30 min in the dark. 12.5 µl of each reaction was loaded on a fresh 6% native acrylamide gel pre-run for at least 30 min in 1X Tris Borate EDTA (TBE) in the dark at 4°C for 1 hr at 100 volts. The gels were imaged using the BioRad Chemidoc MP imaging system with a 30 sec exposure and the manufacturer’s settings for AlexaFluor 488 detection.

For reactions where the binding of RtrB was tested, DNA binding reactions were performed in 20 µl reaction volumes containing binding buffer (20mM HEPES pH 7.8, 4 mM MgCl_2_, 37 mM KCl, 1 mM CaCl_2_, 0.75 mM dithiothreitol, 8% glycerol and 75 µg/ml bovine serum albumin), 7 ng of fluorescently labeled DNA probe and increasing concentrations of RtrB (0, 10, 25, 50, 75, 100, 250 nM). For competition experiments, a 10-fold excess of cold competitor probe was added to the reaction. Reactions were incubated at room temperature for 30 min in the dark. 12.5 µl of each reaction was loaded on a fresh 6% native acrylamide gel pre-run for at least 30 min in 1X TAE in the dark at 4°C for 30 min at 80 volts, and subsequently 1 hr at 100 volts. The gels were imaged using the BioRad Chemidoc MP imaging system with a 30 sec exposure and the manufacturer’s settings for AlexaFluor 488 detection.

### RNA preparation, sequencing and analysis

Strains grown on PYE agar plates were inoculated into 5 ml of M2X medium in 3 biological replicates for each strain, and grown overnight on a rolling incubator at 30°C. Cultures were diluted to OD660nm of 0.008 in 8 ml of fresh M2X and grown in the same manner until they reached OD_660nm_ of 0.30 – 0.35. At this point, 6 ml of each replicate were collected by centrifugation at 15,000 × g for 1 min. This cell pellet was immediately resuspended in 1 ml of TRIzol and stored at −80° C. The samples were heated for 10 min at 65°C. Then, after addition of 200 µl of chloroform samples were vortexed and incubated at room temperature for 5 min. Phases were separated by centrifugation at 15,000 × g for 15 min at 4°C. The upper aqueous phase was transferred to a new tube and 0.7 volume of cold 100% isopropanol was added. Samples were then stored at −80°C overnight. The overnight precipitation was centrifuged at 15,000 × for 30 min at 4°C and the resulting nucleic acid pellet was washed twice with cold 70% ethanol. The pellet was centrifuged again at 15,000 × for 5 min at 4°C, ethanol was removed and the pellet was allowed to dry. The nucleic acid pellet was resuspended with nuclease-free water. The samples were treated with Turbo DNase (Ambion, Life Technologies) and clean-up was performed with RNeasy Mini Kit (Qiagen).

Stranded cDNA libraries were prepared and sequenced by the Functional Genomics Core at the University of Chicago. Briefly, RNA samples were treated with the Ribo-zero Kit (Illumina) for rRNA removal and library preparation proceeded with Illumina ScriptSeq RNA-Seq Library Preparation Kit. All the libraries were sequenced with an Illumina HiSeq4000 instrument. Analysis of RNA-seq data was performed with CLC Genomics Workbench (Qiagen). Reads were mapped to the *C. crescentus* NA1000 genome (Genome accession number CP001340). We used the RNA-seq Analysis Tool to analyze the differential expression between the different strains.

### Data availability

The raw RNA-seq data for each sample are available in the NCBI Gene Expression Omnibus (GEO) under accession number GSE125783.

### SpdR binding motif discovery

We used the MEME Suite [49] to identify potential SpdR binding motifs in promoter regions of genes with more than 2-fold differences in expression between the *spdR*(D64E) and Δ*spdR* strains. The promoter sequences (−150 to +50, based on translation start site annotations in GenBank accession CP001340) of genes with > +2-fold change (up-regulated in *spdR*(D64E) relative to Δ*spdR*) were submitted to MEME to identify motifs. A motif with high similarity to that previously reported for *C. crescentus* SpdR [39, 40] and for SpdR homologs [46–48] was found in 65 of 119 promoter sequences, each with a p-value < e^−4^ (listed in Table S2; see Fig 5). Using the FIMO Motif Scanning algorithm, we applied the SpdR motif matrix from MEME to scan the promoter sequences (−150 to +50) of genes with < −2-fold change (down-regulated by *spdR*(D64E) strain relative to Δ*spdR*) for potential SpdR motif sequences. Predicted SpdR binding motifs with a p-value cutoff < e^−4^ were present in the promoter sequence of 17 out of 48 of the down-regulated genes, and are listed in Table S2.

## Acknowledgements

We thank Olivia Stovicek for assistance building strains. We thank past and present members of the Crosson laboratory for helpful discussions.

## Authors Contributions

Conceived and designed the experiments: LMRR AF SC. Performed the experiments: LMRR AF. Analyzed the data: LMRR AF SC. Wrote the paper: LMRR AF SC.

## Figures

**Fig S1.**
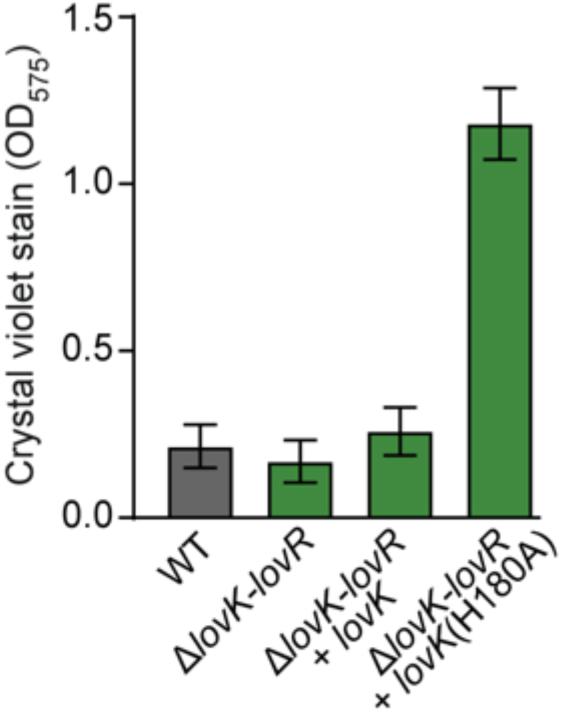
*lovR* is not required for the *lovK*(H180A) regulated increase in surface adhesion. Strains bearing a deletion of the *lovK-lovR* locus were complemented with a wild-type *lovK* or *lovK*(H180A) allele. Surface attachment to polystyrene plates was measured by crystal violet stain. Cultures were grown in M2X medium. Data are representative of at least three independent experiments. Bars represent mean ± s.d.; n=7.

**Fig S2.**
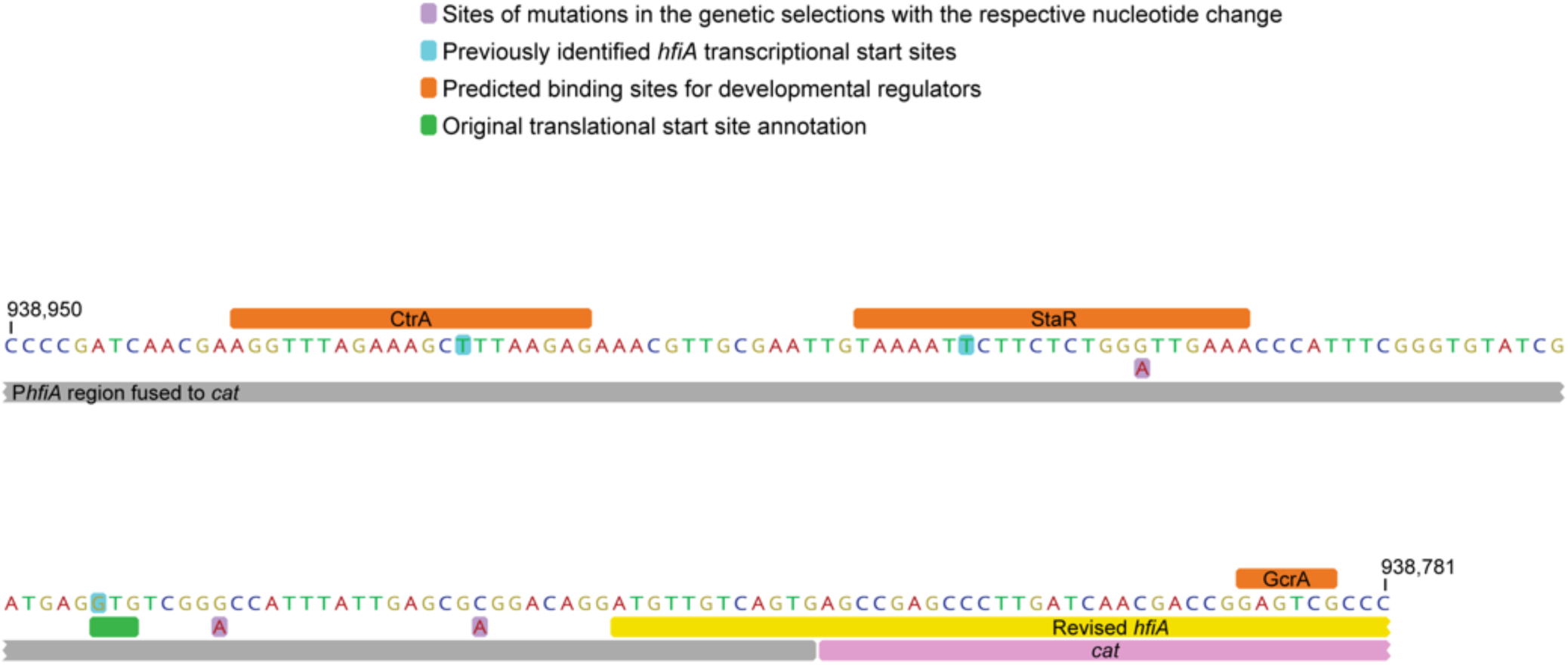
Diagram of the *hfiA* promoter region. Predicted DNA-binding regions of several developmental regulators in orange [14]. Original annotated start codon of *hfiA* in green; the revised *hfiA* reading frame is in yellow [14]. Experimentally identified *hfiA* transcriptional start sites in light blue [14] [73]. The central portion of the P*hfiA-cat* fusion is shown with P*_hfiA_* in gray while and cat in pink. Sites of spontaneous mutations identified in the genetic selections with the respective nucleotide change in light purple. Genome coordinates are based on the NA1000 genome.

**Fig S3.**
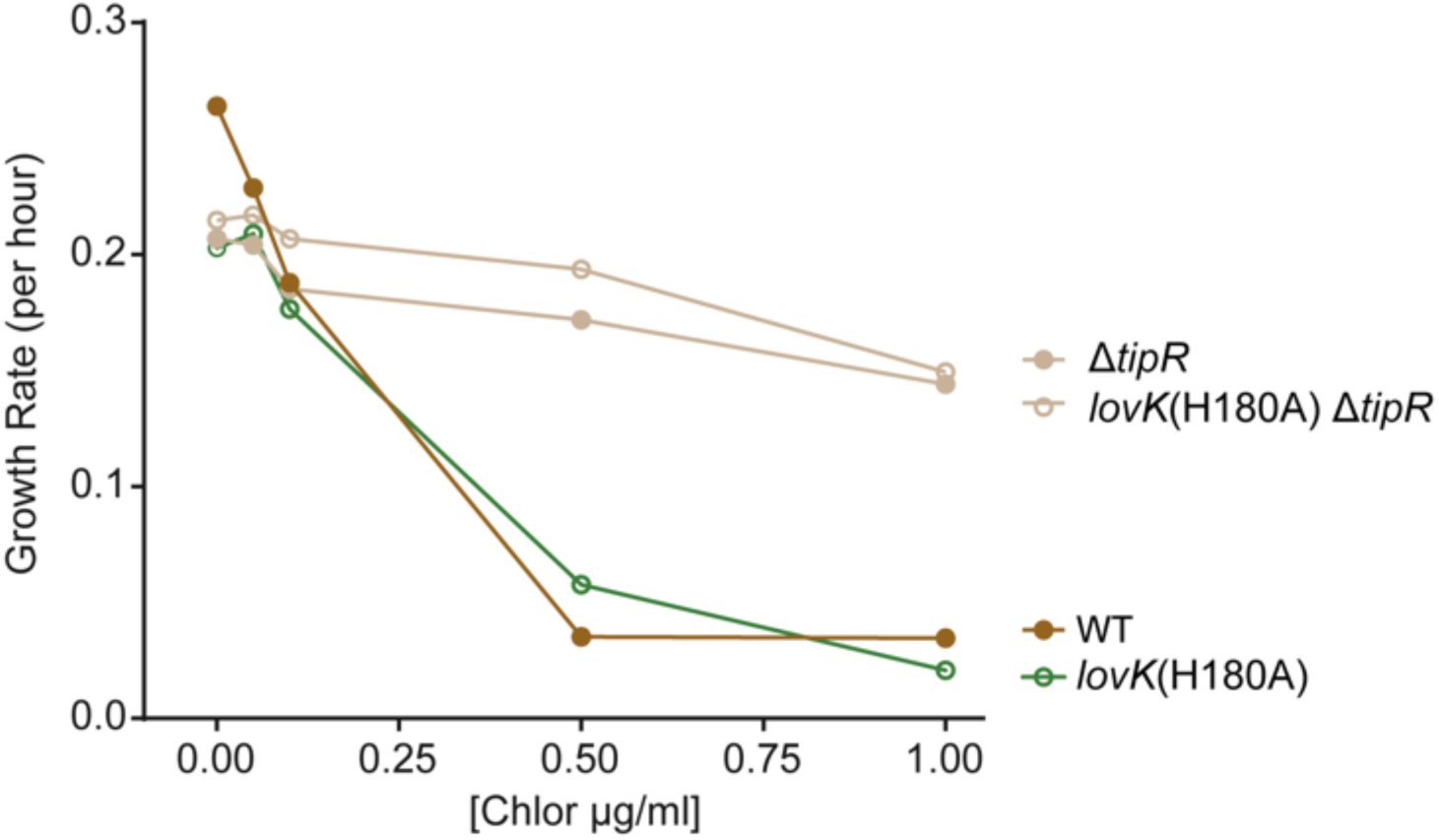
Strains carrying a *tipR* in-frame deletion are resistant to chloramphenicol. Growth rate per hour at different chloramphenicol concentrations for wild-type (WT), *lovK*(H180A), Δ*tipR*, and *lovK*(H180A) Δ*tipR* strains. Cultures were grown in M2X medium supplemented to the final chloramphenicol concentrations indicated on the x-axis. One biological replicate per condition per strain.

**Fig S4.**
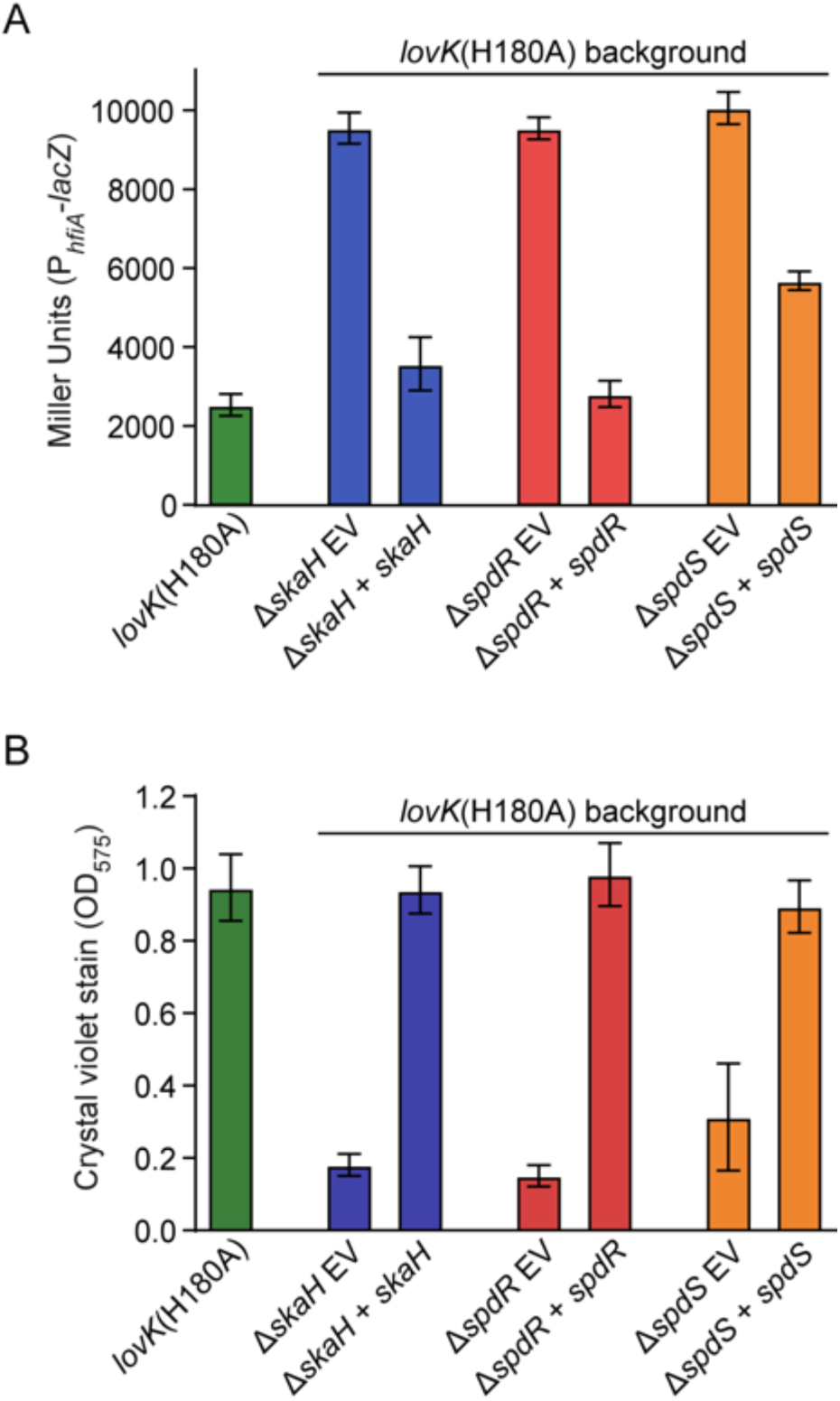
Complementation of the in-frame deletions in a *lovK*(H180A) background. Strains bearing in-frame deletions of *skaH, spdS* or *spdR* in a *lovK*(H180A) background were transformed with either pMT680 plasmids that express the corresponding disrupted gene from a xylose inducible promoter or a pMT680 as an empty vector (EV) control. **A)** β-galactosidase activity from the P*_hfiA_-lacZ* transcriptional fusion was measured in each strain. Cultures were grown in M2X medium to OD_660nm_ of 0.05 - 0.15. Presented data are representative of at least three independent experiments. Bars represent mean ± s.d.; n=6. **B)** Surface attachment of cells grown in polystyrene plates was measured by crystal violet stain. Cultures were grown in M2X medium for 16 hrs post-inoculation. Data are representative of three independent experiments. Bars represent mean ± s.d.; n=9.

**Fig S5.**
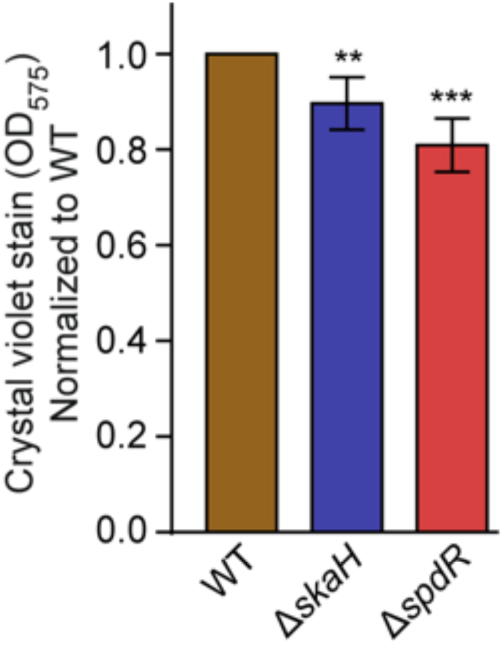
Deletion of *skaH* or *spdR* decreases surface attachment. Crystal violet stain from cells growing in polystyrene plates from seven different experiments was collected. The mean Δ*skaH* and Δ*spdR* crystal violet stain for each experiment was normalized to the mean wild-type stain of the respective day. One-way ANOVA followed by Dunnett’s multiple comparisons test was performed using GraphPad Prism version 8.0.0. to compare wild-type to the mutant strains*. ** P<*0.005, *** *P*<0.0005

**Fig S6.**
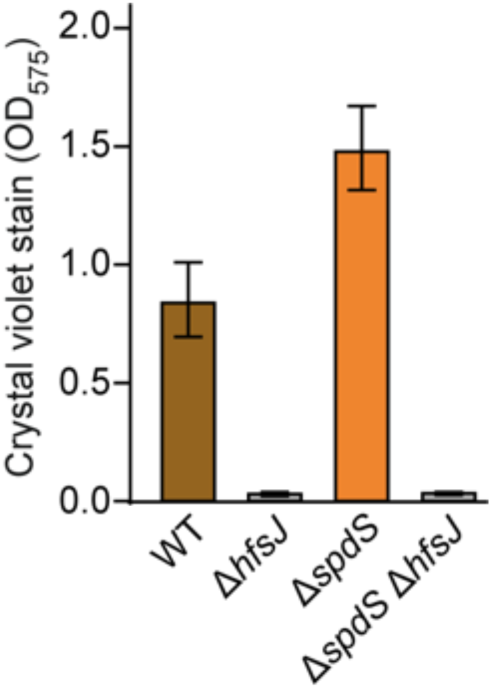
Holdfast synthesis is required for the Δ*spdS* stationary phase hyper-attachment phenotype. Surface attachment to polystyrene plates was measured by crystal violet stain in wild-type (WT), Δ*spdS*, Δ*hfsJ* and Δ*spdS* Δ*hfsJ* strains. Cultures were grown in M2X medium until stationary phase (24 hrs). Bars represent mean ± s.d.; n=12.

**Fig S7.**
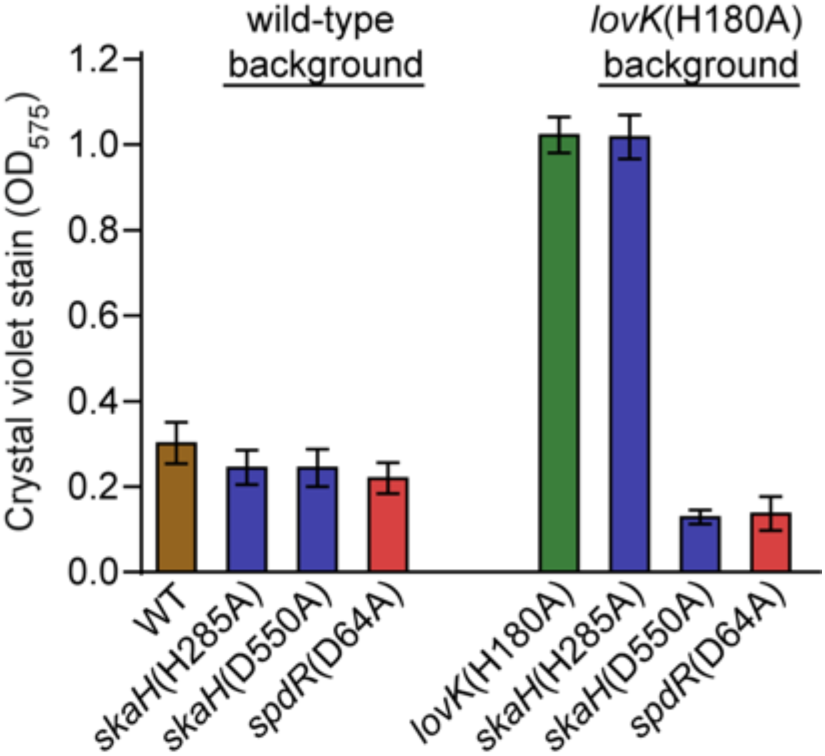
Surface attachment of strains bearing mutations in the TCS protein phosphorylation sites. Strains bearing point mutations in the conserved phosphorylation sites of *skaH, spdS* or *spdR* in either a wild-type (WT) or *lovK*(H180A) background were grown in M2X medium in polystyrene plates for 16 hrs post inoculation. Surface attachment was measured by crystal violet stain. Data presented are representative of three independent experiments. Bars represent mean ± s.d.; n=8.

**Fig S8.**
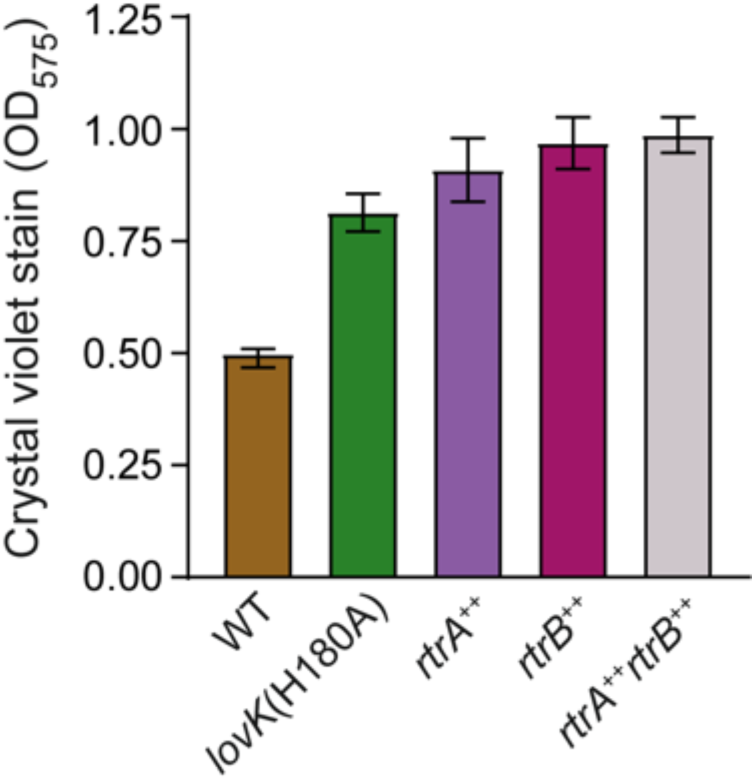
Overexpressing the transcription factors *rtrA* and *rtrB* increases surface adhesion. Surface attachment of cells grown in polystyrene plates was measured by crystal violet stain. Wild-type (WT) and *lovK*(H180A) carry empty vectors of pMT680 and pMT585. *rtrA*^++^ carries pMT680-*rtrA* and pMT585 empty vector. *rtrB*^++^ carries pMT585-*rtrB* and pMT680 empty vector. *rtrA*^++^*rtrB*^++^ carries pMT680-*rtrA* and pMT585-*rtrB*. Cultures were grown for 16 hrs post-inoculation in M2X medium. Data presented are representative of at least three independent experiments. Data represent mean ± s.d.; n=6-8.

**Fig S9.**
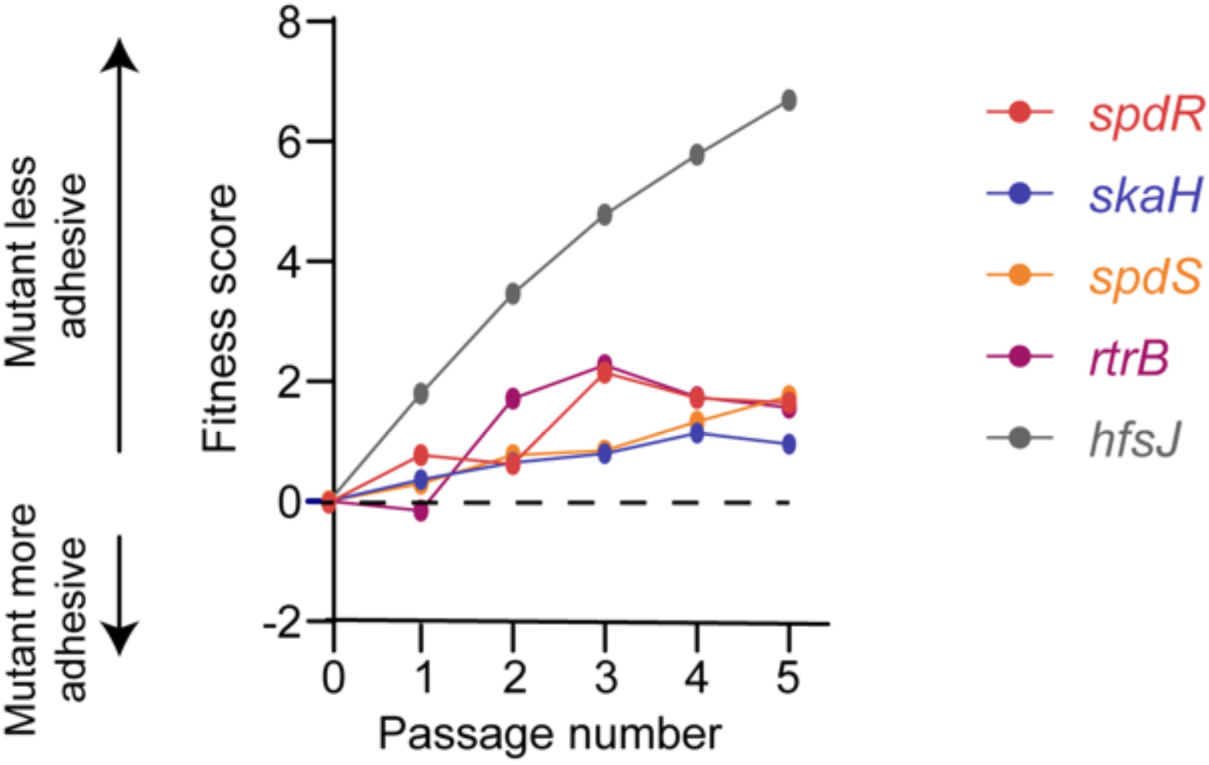
Global Tn-seq approach provides an additional line of evidence that genes identified in this study play a role in regulating adhesion in wild-type cells. Extracted fitness profiles of mutants characterized in a genome-wide attachment screen [58] Briefly, a *C. crescentus* transposon mutant library was grown and passaged in complex PYE medium in the presence of cheesecloth to provide surface area for cell attachment. Cells were periodically sampled from the broth over several days. Hyper-adhesive mutants are titrated out of the broth by the cheesecloth and, consequently, have negative fitness scores relative to the average strain. Conversely, hypo-adhesive mutants are enriched in the broth resulting in positive fitness scores. Strains bearing transposon insertions in TCS genes identified in our study have positive fitness scores. *hfsJ* mutants, which do not produce holdfast and are non-adhesive, are presented as a reference.

**Table S1. Genetic lesions in mutants identified in the chloramphenicol growth selections**

**Table S2. Transcripts with more than 2-fold difference between *spdR*(D64E) and Δ*spdR***

**Table S3. Plasmids, primers and strains used**

**Table S4. RNA-seq analysis for all genes**

